# Single-cell transcriptomic analysis of NK cell dynamics in myeloma patients reveal persistent reduction of cytotoxic NK cells from diagnosis to relapse

**DOI:** 10.1101/2023.07.05.547295

**Authors:** Sabrin Tahri, Madelon M.E. de Jong, Cathelijne Fokkema, Natalie Papazian, Zoltán Kellermayer, Michael Vermeulen, Mark van Duin, Gregory van Beek, Remco Hoogenboezem, Pieter van de Woestijne, Kazem Nasserinejad, Elona Saraci, Mattia D’Agostino, Francesca Gay, Vincent H.J. van der Velden, Mathijs A. Sanders, Sonja Zweegman, Niels W.J.C. van de Donk, Annemiek Broijl, Pieter Sonneveld, Tom Cupedo

## Abstract

Natural killer (NK) cells mediate the cytotoxic immune response against multiple myeloma and are important effector cells in immune therapies through antibody-dependent cellular cytotoxicity. Here, we used single-cell transcriptomics, flow cytometry and functional assays to investigate the bone marrow NK cell compartment of myeloma patients at diagnosis, during treatment and after relapse. The bone marrow of myeloma patients is characterized by a reduction in conventional cytotoxic NK cells that persists throughout treatment. We show in 20% of newly diagnosed myeloma patients that an altered balance between cytotoxic and cytokine-producing NK cells translates into a reduced cytotoxic ability in response to therapeutic antibodies. The relative loss of cytotoxic NK cells persists at relapse and is accompanied by an expansion of IFN-responsive NK cells. These findings reveal previously unappreciated alterations in bone marrow NK cell composition and highlight the importance of understanding the bone marrow immune system in patients receiving immunotherapies.

**Statement of significance:** The bone marrow of multiple myeloma patients is characterized by a persistent reduction in cytotoxic CD56^dim^ NK cells, accompanied by inferior *in vitro* responses to therapeutic antibodies at diagnosis and an increase in IFN-responsive NK cells at relapse. These findings highlight the importance of understanding the BM microenvironment in multiple myeloma patients receiving immunotherapies.

## Introduction

Despite improved understanding of disease pathophysiology and development of novel therapies, multiple myeloma (MM) remains incurable^1^. MM disease progression and therapy response result from tumor cell-intrinsic features and tumor cell-extrinsic cues from the bone marrow (BM) microenvironment^2^. Natural killer (NK) cells are important effector cells of the cytotoxic immune response against MM^3^, and efficacy of existing MM therapies, including immunomodulatory drugs and monoclonal antibodies, relies substantially on the function of endogenous NK cells^4^.

NK cells are innate lymphoid cells (ILCs) that exert cytotoxicity against stressed cells, including malignant (plasma) cells^3, 5^. NK cells are divided into a cytotoxic CD56^dim^ subset, important for antibody-depended cellular cytotoxicity (ADCC), and a cytokine-producing CD56^bright^ subset which secrete inflammatory mediators such as interferon (IFN)-γ, tumor necrosis factor (TNF)-α and granulocyte-macrophage colony-stimulating factor (GM-CSF)^6^. Alterations in NK cell functionality and subset distribution are associated with inferior clinical outcomes in solid malignancies^7, 8, 9, 10^. Single-cell transcriptomic analyses demonstrated similar alterations in BM NK cell subsets, with repressed NK cell effector functions accompanied by heterogeneity in the frequencies of CD56^bright^ and CD56^dim^ subsets in AML patients^11^.

In MM patients, NK cells have lower expression of activating receptors NKG2D^12,13^, CD16, 2B4^14^, and DNAM-1^15^, and increased expression of inhibitory receptors KIR2DL1^16^ and PD1^17, 18^. Moreover, peripheral blood NK cells exhibit reduced cytotoxicity against MM cells *ex vivo*^3^. These alterations suggest that the immunosuppressive tumor niche induces dysfunctional NK cells^4, 19^, likely impacting anti-MM NK cell cytotoxicity and efficacy of therapies dependent on ADCC, such as anti-CD38 monoclonal antibodies. Indeed, while CD38-targeting monoclonal antibodies are becoming the global standard of care for newly diagnosed MM (NDMM) patients^20^, variability in depth of response among patients exists^21^. Further efforts to predict and improve individual responses to therapeutic antibodies in MM are needed.

Here, we set out to investigate the composition and function of the BM NK cell compartment in MM patients at diagnosis, during treatment, and after relapse using single-cell transcriptomics, flow cytometry and functional assays. Comparison of transcriptomic profiles of BM NK cells from NDMM patients with non-cancer controls revealed reduced frequencies of cytotoxic NK cells in MM. In approximately 20% of NDMM patients, this reduced cytotoxic potential translated into inferior responses to therapeutic antibodies in *in vitro* functional assays. The reduction in cytotoxic NK cells persisted during anti-myeloma therapy and, after relapse, was accompanied by an expansion of IFN-responsive NK cells. These findings reveal substantial dysregulation of the BM NK cell compartment with potential implications for therapies dependent on ADCC. Our study offers a temporal overview of the NK cell compartment across therapy and disease progression, providing a rich resource for analyses of an important effector cell within the MM BM microenvironment.

## Results

### Single cell transcriptomic landscape of BM NK cells

To define the BM NK cell landscape in MM patients, we generated a single-cell transcriptomic dataset of 57,822 CD38^+^BM NK cells (9,146 cells from controls and 48,315 cells from MM patients; Figures 1A-C; see Methods, Supplemental Figures 1A-B)^22^. Non-paired samples were collected at diagnosis (n=19), after induction therapy (cycle 4 day 28; C4D28, n= 3), after consolidation therapy (day 100; D100, n=6), during maintenance therapy (week 25, week 52, week 105; W25, W52, W105, n=12) and at disease relapse (n=6) from transplant-eligible NDMM patients treated with bortezomib, thalidomide and dexamethasone with or without daratumumab (HOVON-131/IFM 2015-01 trial; Supplemental Table 1, Supplemental Figure 2)^23^. After pre-processing and quality control (see Supplemental Methods), UMAP dimensionality reduction identified 7 NK cell clusters (Figures 1B-C).

**Figure 1.**
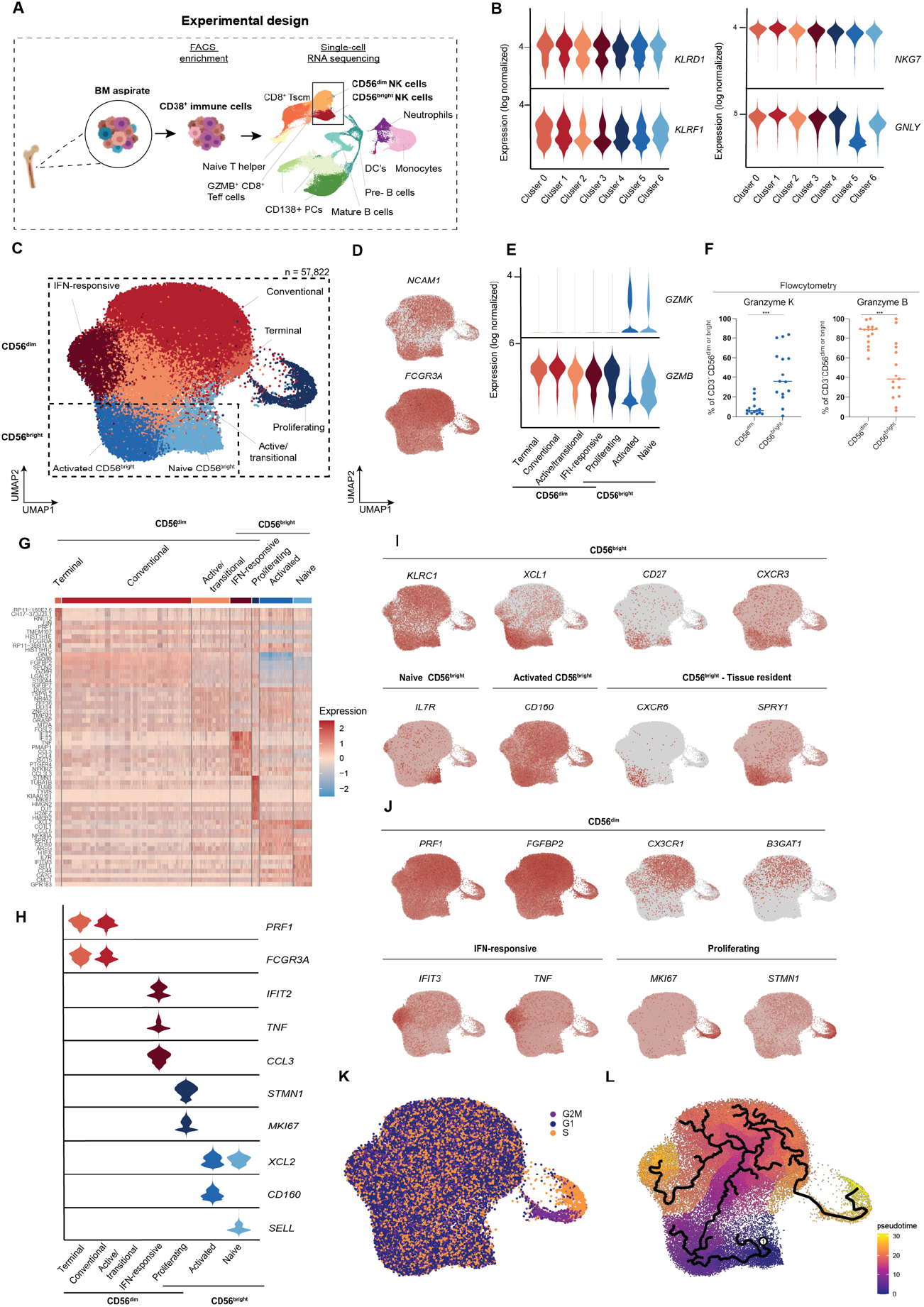
Single-cell transcriptomic landscape of BM NK cells in MM. (A) Experimental design depicting FACS enrichment of CD38^+^ immune cells from BM aspirate followed by single-cell RNA sequencing. (B) Violin plots demonstrating the transcription of four NK-lineage defining markers per cluster. (C) UMAP of 57,822 human CD38^+^BM NK cells from MM patients and controls identifying seven distinct clusters. Cells are color-coded according to the defined clusters. (D) Feature plot depicting *NCAM1* (CD56) and *FCGR3A* (CD16) transcription among NK cell clusters. (E) Violin plots demonstrating *GZMK* and *GZMB* transcription among the 7 clusters of NK cells. (F) Flow cytometric analyses of Granzyme K and Granzyme B protein in CD56^bright^ and CD56^dim^ NK cells from MM BM (n=15). (G) Heatmap of the top differentially expressed genes in depicted cluster compared to all other clusters. (H) Violin plots of marker genes characterizing the seven clusters depicted in figure 1C. (I) Transcription of marker genes characterizing CD56^bright^ NK cell clusters. (J) Transcription of marker genes characterizing CD56^dim^ NK cell clusters. (K) Transcription of cell cycle associated genes in the CD38^+^ NK cell dataset. (L) Monocle 3 pseudotime analysis of NK cells. ‘1’ represents the selected starting point of the trajectory analysis. The data in (F) is depicted as median. Significance was calculated using the Mann-Whitney U test (two-tailed); NS, P>0.05; **P≤0.01.

Since NK cells are functionally divided in cytotoxic CD56^bright^ and cytokine-producing CD56^dim^ NK cells^6^, we assigned clusters to these major subsets based on transcriptional features. *NCAM1* transcripts (encoding CD56) are insufficiently discriminatory in single-cell transcriptomic datasets to discern CD56^dim^ and CD56^bright^ NK cells^24^ (Figure 1D), and we therefore used transcription of *GZMB* versus *GZMK* to identify the two subsets^25^ (Figure 1D-E). Differential enrichment of Granzyme B or Granzyme K protein was verified by flow cytometry (Figure 1F, gating strategy in Supplemental Figure 1C), and further confirmed by *FCGR3A* (encoding CD16) transcription (Figure 1D). As a result, cluster 0, 1 and 2 were defined as cytotoxic *GZMB*^+^CD56^dim^ NK cells, cluster 5 and 6 as *GZMK*^+^CD56^bright^ NK cells, and clusters 3 and 4 contained a mixture of these two NK cell subsets (Figure 1E).

Next, we annotated the individual clusters within these CD56^bright^ and CD56^dim^ NK cell compartments by differentially expressed genes and canonical and previously reported markers of NK cell subtypes (Figures 1G-J, Supplemental Figure 1D)^11, 26, 27^. Overall, the clusters of *GZMK*^+^CD56^bright^ NK cells had increased transcription of *KLRC1, XCL1, XCL2, CD27* and *CXCR3*, while *GZMB*^+^CD56^dim^ clusters were characterized by *CX3CR1, B3GAT1, KIR2DL1, KIR2DL3, LGALS1, S1PR5* and several cytotoxicity markers (*PRF1, FGFBP2, SPON2* and *GNLY*) (Figures 1G-I, Supplemental Figure 1D). CD56^bright^ NK cells included a naïve *SELL*^+^*IL7R*^+^ cluster and an activated *CD160*^*+*^*TOX2*^+^ cluster (Figures 1G-I, Supplemental Figure 1D). The latter additionally transcribed tissue-residency markers such as *CXCR6, CCR5* and *SPRY1/2* (Figure 1I), and had transcriptomic overlap with recently described lymphoid tissue-resident (lt) NK cells^27^ and hNK_Bm2^11^.

The CD56^dim^ compartment contained 3 uniformly *GZMB*^+^ clusters and 2 clusters that additionally had up to 20% *GZMK*^+^ NK cells (Figures 1G-H, J, Supplemental Figure 1D). The conventional cytotoxic CD56^dim^ cluster was characterized by cytotoxicity markers (*PRF1, FGFBP2, NKG7, GNLY, CST7, PRF1* and *GZMB*) and *CX3CR1*, while the transitional/recently activated NK cluster combined intermediate expression of these genes with the expression of immediate early genes (e.g. *DUSP2, NR4A2* and *FOSL2*) (Figure 1G, Supplemental Figure 1D)^11, 26, 27, 28^. The terminal NK cluster was transcriptionally similar to the conventional cytotoxic CD56^dim^ cluster but had increased average expression of several cytotoxicity genes including *FCGR3A* and *PRF1* (Supplemental Figures 1D-E). In line with other NK cell datasets^26, 27, 28^, transcription of CD57 (*B3GAT1*), the functionally distinguishing marker for terminally differentiated NK cells, was too low to differentiate conventional cytotoxic NK cells from terminal NK cells (Supplemental Figure 1E). The same was true for other maturation markers including *ZEB2, CX3CR1* and *HAVCR2* (Supplemental Figure 1E). Instead, the terminal NK cluster was identified by increased transcription of histone subunits (*HIST1H1E, HIST1H1C, HIST1H2AC, HIST1H1D* and *HIST2H2BF*), small nucleolar RNA and DNA repair and damage response genes (*RNU12, EGR1, NEAT1*)^26^, genes related to cytoskeleton remodeling (*ACTB, PFN1, CORO1A, CFL1, CLIC1* and *ACTR3*), and by a relative decrease of ribosome-related genes (Supplemental Figures 1E-G). The 2 *GZMB*^*+*^ clusters with *GZMK*^+^ NK cells (Supplemental Figure 1H) contained an IFN-responsive cluster defined by increased expression of interferon-related genes and immediate early genes, such as *IFIT3, ISG15, FOSB* and *NR4A2*, and *TNF* (Figure 1J, Supplemental Figure 1D) and a cluster of proliferating NK cells identified by enrichment for proliferation and cell-cycle related genes such as *MKI67* and *STMN1* (Figure 1J) ^27^. Cell cycle analysis confirmed increased expression of G2M and S phase transcripts in this latter cluster (Figure 1K). The developmental trajectory of MM BM NK cells was mapped by pseudotime analysis and revealed, when assigning the naïve *GZMK*^+^CD56^bright^ NK cell cluster as starting point, that trajectories progress via the transitional/active NK cell cluster and conventional cytotoxic CD56^dim^ NK cells into the direction of either the IFN-responsive or terminal and proliferating *GZMB*^*+*^CD56^dim^ clusters (Figure 1L). This validated the existence of a transitional NK cell cluster in MM BM, in line with previous studies ^11, 26, 27, 28^. Calculation of a module score based on the gene expression of helper ILC ^29, 30^ did not identify helper ILCs in our dataset (Supplemental Figure 1I), which is likely due the exclusion of *NKG7* low cells during in silico pre-processing^31^.

### Reduction of cytotoxic CD56^dim^ NK cells and an increase of activated CD56^bright^ NK cells in NDMM

To identify BM NK cell alterations in NDMM, we first compared MM patients at diagnosis with controls. The previously identified 7 BM NK cell clusters were present in all NDMM patients and all controls (Figure 2A), and analysis of differentially expressed genes in the individual clusters from NDMM vs controls revealed no major differences (Supplemental Table 2). This also translated in an absence of transcriptionally enriched gene pathways in individual clusters of NDMM NK cells (data not shown). Nonetheless, we did detect differences in the abundance of clusters when comparing NDMM to controls. The conventional cytotoxic CD56^dim^ NK cell cluster was significantly reduced in NDMM BM, while there was a significant increase in activated CD56^bright^ NK cells (Figure 2B). This altered balance in major CD56^dim^ and CD56^bright^ clusters was reflected in the genes differentially expressed in the total BM NK cell pool of NDMM patients versus controls. In NDMM patients, the total BM NK cell pool revealed a reduction in cytotoxicity genes and an increase in IFN-response genes (Figure 2C), suggesting a shift away from cytotoxicity. To investigate the extent of this reduced cytotoxic potential we analyzed key transcriptional features associated with cytotoxicity and general NK cell function. BM NK cells in NDMM patients had reduced abundance of cytotoxicity genes (e.g. *NKG7, GNLY* and *PRF1*), increased abundance of inhibitory receptors *KLRB1* (CD161) and *KLRC1* (NKG2A), and decreased presence of activating receptors *NCR3* (NKp30), *CD226* (DNAM-1) and *KLRK1* (NKG2D) (Figure 2D). The reduced abundance of CD56^dim^ NK cells in NDMM patients was accompanied by a reciprocal increase in cytokine-producing CD56^bright^ NK cells, which was reflected in an increase of pro-inflammatory cytokines and chemokines including *CCL3* (MIP-1α), *XCL1/2, IL6ST, IFNG* and *TNF* (Figure 2D). Combined, these transcriptomic alterations suggest an imbalance in *GZMB*^+^CD56^dim^ vs. *GZMK*^+^CD56^bright^ NK cells in NDMM patients that leads to reduced cytotoxic potential and an increased pro-inflammatory signature of the total BM NK cell pool.

**Figure 2.**
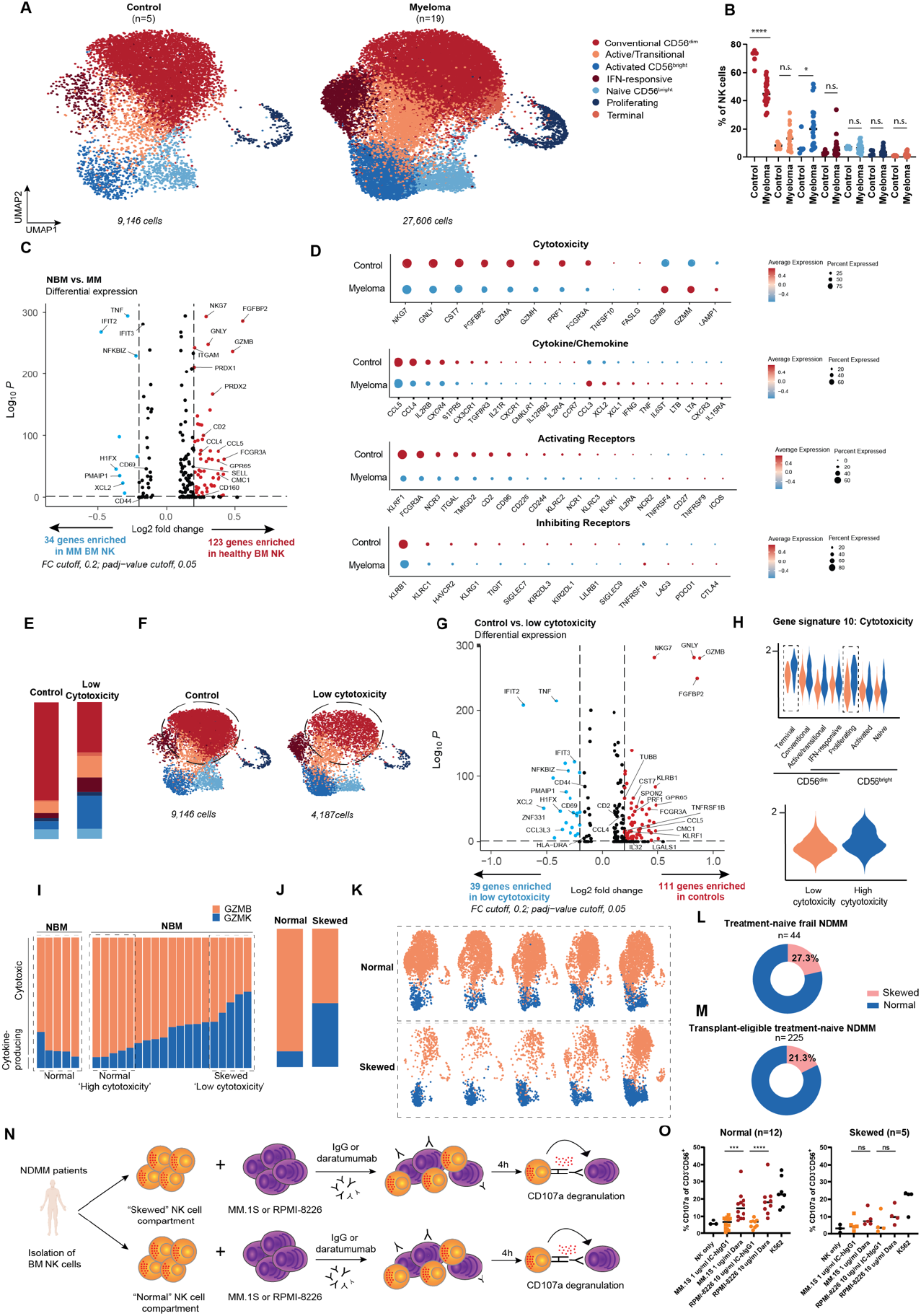
NDMM patients have reduced frequencies of cytotoxic CD56^dim^ NK cells with inferior cytotoxic responses to therapeutic antibodies *in vitro*. (A) Original UMAP as depicted in figure 1C, split by controls (n=5) vs NDMM patients (n=19). (B) Scatter plot depicting frequency of NK cell clusters, split by non-cancer controls vs NDMM patients. (C) Volcano plot depicting differentially expressed genes in the NDMM dataset vs the control dataset; log2 fold change (FC) cutoff, 0.02; adjusted p-value, 0.05. (D) Dot plot depicting transcription of cytotoxicity genes (upper left), cytokine/chemokine genes (upper right), NK cell activating (lower left) and inhibiting receptor (lower right) genes in control vs myeloma NK cells. (E) Bar plot depicting NK cell cluster composition in controls (n=5) vs NDMM patients with lowest frequency of conventional cytotoxic CD56^dim^ NK cells, depicted as ‘low cytotoxicity’ (n=5). (F) Original UMAP as depicted in figure 1C, split by controls (n=5) vs NDMM patients with lowest frequency of conventional cytotoxic CD56^dim^ NK cells (low cytotoxicity, n=5). (G) Volcano plot depicting differentially expressed genes in NDMM patients with lowest frequency of conventional cytotoxic CD56^dim^ NK cells (low cytotoxicity) vs controls; log2 fold change (FC) cutoff, 0.02; adjusted p-value, 0.05. (H) Cytotoxicity gene signature as identified by NMF depicted for NDMM patients with highest or lowest frequency of conventional cytotoxic CD56^dim^ NK cells (high cytotoxicity, n=5 vs low cytotoxicity, n=5). (I) Bar plot depicting cytokine-producing *GZMK*^+^CD56^bright^ NK cells vs cytotoxic *GZMB*^+^CD56^dim^ NK cell composition per individual. Relative loss of cytotoxic NK cells is labelled as skewed. (J) Bar plot depicting cytokine-producing *GZMK*^+^CD56^bright^ NK cells vs cytotoxic *GZMB*^+^CD56^dim^ NK cell composition in NDMM patients with skewed (n=5) or NDMM patients with normal NK cell compartment (n=5). (K) Original UMAP as depicted in figure 1C split by NDMM patients with most skewed (n=5) or normal NK cell composition in the BM (n=5). (L) Donut chart depicting frequency of relative loss of cytotoxic NK cells (‘skewed’) in treatment-naïve frail NDMM patients (n=44). (M) Donut chart depicting frequency of relative loss of cytotoxic NK cells (‘skewed’) in transplant-eligible treatment-naïve NDMM patients (n=225). (N) Experimental design depicting the analysis of daratumumab-mediated CD107a degranulation in response to a co-culture with MM.1S or RPMI-8226 after treatment with IgG1 or daratumumab using BM NK cells from NDMM patients with skewed or normal NK cell compartment by flowcytometry. (O) Dot plots showing percentages of CD107a-positive NK cells, from patients with a normal or skewed NK cell compartment, in response to co-culture with MM cell lines in the presence of IgG or daratumumab. Significance was calculated using the Mann-Whitney U test (two-tailed); NS, P>0.05; *P≤0.05; **P≤0.01; ***P≤0.001; ****P≤0.0001.

The reduced cytotoxic potential of the BM NK cell pool in NDMM patients was further supported by non-matrix factorization (NMF)^32^, a method which extracts gene expression signatures reflective of cell identities or active gene expression programs, even when present in a small cellular subsets or across multiple clusters^33^. Using this method, an activation score is assigned to each cell for each extracted gene signature dependent on the activity of the associated transcriptional program, making a direct comparison between cells possible. NMF identified 15 gene signatures across all NDMM NK cells in our cohort (Supplemental Table 3). A number of NMF signatures confirmed the Louvain algorithm-based cluster identification from Seurat, i.e. IFN-responsiveness (signature 4), activated CD56^bright^ status (signature 7) and proliferation (signature 14; Supplemental Figures 3A-C, Supplemental Table 3). Newly discovered signatures represented functional states such as several activation states, but we also discovered gene signatures related to cytotoxicity, CD56^dim^ status and chemokine profile. Interestingly, 3 NMF signatures were found to be differentially expressed in NDMM compared to controls. Two of these were related to activation (signature 3: *DUSP2, ZFP36* and signature 6: *JUN, FOS*) and were increased in MM patients, while another cytotoxicity-related signature was significantly decreased in NDMM patients (Supplemental Figures 3D-F). NMF confirms the reduced cytotoxic potential of the BM NK cell pool in NDMM patients and identifies 2 transcriptional activation states specifically increased in the conventional cytotoxic CD56^dim^ NK cells of MM patients (Supplemental Figures 3D-F).

### Heterogeneity of NK cell cluster distribution in NDMM patients

Even though the conventional cytotoxic CD56^dim^ cluster was reduced in all NDMM patients compared to controls, the degree of reduction varied between patients (Figure 2B, Supplemental Figure 4A). We hypothesized that the largest differences in functionality of the BM NK cell pool would be found in the NDMM patients with the most significant reduction of the conventional cytotoxic CD56^dim^ cluster. To test this hypothesis, we compared the 5 patients with the lowest percentages of the conventional cytotoxic CD56^dim^ cluster (depicted as ‘low cytotoxicity’ in Figure 2E-G) to healthy controls (Figures 2E-F). This revealed significant reduction in genes associated with cytotoxicity and increase in inflammatory/activation genes in NDMM (Figure 2G, Supplemental Table 4). In line with these findings, NMF analysis revealed that MM patients with the lowest abundance of conventional cytotoxic CD56^dim^ NK cells (depicted as ‘low cytotoxicity’ in Supplemental Figure 3) had a concomitant decrease in the CD56^dim^ gene signature, the cytotoxicity gene signature, and the less-well characterized *GNLY* high and *CCL5* high gene signatures, and an increase of the proliferating gene signature in proliferating NK cells (Figure 2H, Supplemental Figure 3G-K). These analyses suggest that functional differences are most obvious in the patients with the strongest reduction in the conventional CD56^dim^ cluster.

### Altered frequencies of CD56^dim^ and CD56^bright^ NK cells in NDMM

The conventional CD56^dim^ cluster is a major cluster of CD56^dim^ NK cells. This raised the possibility that the reduction in this cluster might be reflected in the CD56^dim^ to CD56^bright^ ratio as traditionally determined by flow cytometry. To explore this possibility, we first evaluated the distribution of total CD56^dim^ versus total CD56^bright^ cells in our single-cell transcriptomics dataset (Figures 2I-K). A subset of NDMM patients had an NK cell distribution that was very similar to controls (those with ‘high cytotoxicity’, hereafter depicted as ‘normal’), while there was a subgroup of patients with a skewed NK cell repertoire caused by a relative loss of cytotoxic cells and a gain of cytokine-producing CD56^bright^ NK cells (hereafter depicted as ‘skewed’). Fittingly, NDMM patients with a skewed NK cell repertoire also have the lowest abundance of conventional cytotoxic CD56^dim^ NK cluster.

To analyze whether these differences would also be apparent at the cellular level, and in which frequency, we analyzed percentages of cytokine-producing CD56^bright^ and cytotoxic CD56^dim^ NK cells in BM of NDMM patients by flow cytometry. The standardized EuroFlow protocol enables reliable minimal residual disease (MRD) detection in MM by two eight-color antibody panels with CD56 as NK cell marker^34^. Of note, this panel was not designed to enumerate NK cells and might still contain dying cells and contaminating CD56-expressing T cells. Nonetheless, we confirmed that we could generate a CD56^bright^ and CD56^dim^ classification by comparing this EuroFlow panel in BM aspirates from NDMM patients with a panel specifically designed for NK cell identification (Supplemental Figures 4B-C). Since NK cell composition in the PB was not reflective of that in the BM (Supplemental Figure 4D), we focused on NK cell subset distribution in the BM only. Typically, normal BM consists of ∼10% CD56^bright^ NK cells and ∼90% CD56^dim^ NK cells^25,35^. Flow cytometric analyses of 269 NDMM patients from 2 independent clinical cohorts revealed reduced frequencies of cytotoxic CD56^dim^ NK cells (“skewed”), defined as <90% CD56^dim^ NK cells, in 27.3% of treatment-naïve frail transplant-ineligible MM patients (n=44, Figure 2L, Supplemental Figure 4E) and in 21.3 % of treatment-naïve transplant-eligible NDMM patients (n=225, Figure 2M, Supplemental Figure 4F).

### Functional consequences of reduction in CD56^dim^ BM NK cells

To investigate whether the most skewed NK cell distribution translated into a reduced functional ability to induce ADCC, we compared daratumumab-induced degranulation of NK cells from patients with skewed and normal NK cell distribution (Figure 2N depicts experimental design). As BM samples were not readily available from healthy controls, NDMM patients with ≥ 90% CD56^dim^ BM NK cells were used as a comparison. In contrast to patients with a normal NK cell distribution, NK cells from patients with a skewed NK cell compartment did not increase degranulation after their co-culture with daratumumab-labeled MM cell lines (Figure 2O, Supplemental Figure 4G), indicating that the variation in BM NK cell composition in NDMM patients can have direct functional consequences on the ability of the BM NK cell response to therapeutic antibodies.

### BM NK cell alterations during treatment

Treatment can severely impact the tumor microenvironment. To assess how MM treatment affects the altered BM NK cell landscape in NDMM we analyzed this compartment throughout treatment and disease relapse. Datasets were generated from 19,009 NK cells from 27 patients during post-induction, post-consolidation and maintenance therapy, as well as at relapse (Figure 3A-B, therapy regimen and clinical trial design summarized in Supplemental Figure 2)^36^. Previous studies described a decrease of total and CD38^+^ NK cells after daratumumab treatment^18^. So to understand the effects of anti-CD38 antibody-containing myeloma treatment on BM NK cells, we first analyzed NK cell composition in patients who were treated with or without daratumumab during induction and consolidation therapy or during maintenance (see Supplemental Figure 2 for trial design) compared to patients who did not receive daratumumab. Daratumumab treatment at any time point resulted in an overall decrease of the cytotoxic *GZMB*^+^ CD56^dim^ NK cells (Supplemental Figure 5A), which fittingly have the highest surface expression of CD38^18, 37^. This concomitantly led to an enrichment of activated CD56^bright^ NK cells prior to daratumumab maintenance and an increase in active/transitional NK cells during daratumumab maintenance (Supplemental Figure 5A).

**Figure 3.**
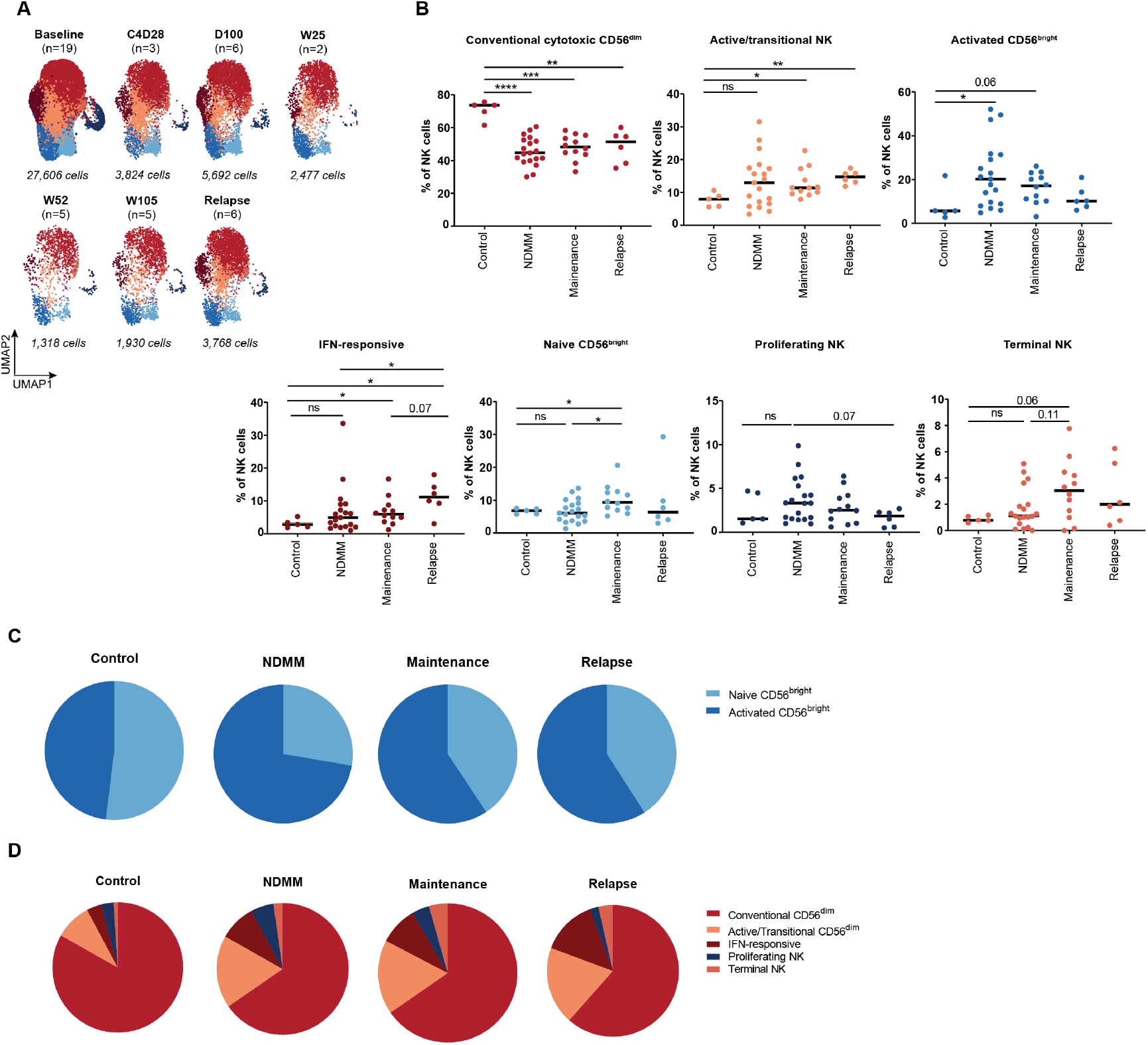
Dynamic alteration in BM NK cells during maintenance and at relapse. (A) UMAPs for MM patients only, split per timepoint. (B) Scatter plots depicting frequency of NK cell clusters per timepoint, split by NK cell cluster. (C) Pie charts depicting frequency of naïve and activated CD56^bright^ NK cell cluster within the CD56^bright^ NK cell compartment, split per timepoint. (D) Pie charts depicting frequency of conventional cytotoxic CD56^dim^ NK cells, active/transitional NK cells, IFN-responsive NK cells, proliferating NK cells and terminal NK cells within the CD56^dim^ NK cell compartment, split per timepoint. Significance was calculated using the Mann-Whitney U test (two-tailed); NS, P>0.05; *P≤0.05; **P≤0.01; ***P≤0.001; ****P≤0.0001.

Next, we generated a longitudinal overview of the BM NK cell compartment. All 7 previously identified NK cell clusters were present throughout intensive therapy, maintenance therapy, and relapse. The decrease in abundance of conventional cytotoxic NK cell cluster, found at diagnosis, persisted throughout treatment, including at relapse (Figure 3B). Compared to diagnosis, naïve CD56^bright^ NK cells increased during maintenance therapy (Figure 3B, Supplemental Figure 5B), reaching normal proportions, but were once again replaced by activated CD56^bright^ NK cells in patients that experienced disease relapse (Figure 3C, Supplemental Figure 5C). Additionally, IFN-responsive CD56^dim^ NK cells progressively increased from diagnosis to relapse (Figure 3D, Supplemental Figure 5D). In sum, a persistent loss of conventional cytotoxic CD56^dim^ NK cells is seen post-therapy, which at relapse is accompanied by expansion of IFN-responsive NK cells.

## Discussion

NK cells are increasingly appreciated for their therapeutic potential in MM, exemplified by monoclonal antibodies directed towards CD38 that have revolutionized the therapeutic landscape^20^. Endogenous BM NK cells are considered primary mediators of the anti-tumor response, yet several studies identified NK cell dysfunctionality at time of myeloma diagnosis^4^. In contrast, the dynamics of NK cell phenotypes in patients undergoing first-line treatment have remained under-explored. Here, using single-cell transcriptomics and validation by flow cytometry in 2 independent cohorts, we show that MM patients have a relative loss of conventional cytotoxic CD56^dim^ NK cells which persists during treatment and disease relapse. Moreover, in approximately 20% of NDMM patients this altered NK cell distribution translates into reduced cytotoxic potential and response to therapeutic antibodies in *ex vivo* assays. Our data identify NK cell alterations at diagnosis and suggests that correcting reduced cytotoxicity could allow for improved therapy responses in the context of therapeutic antibodies.

Differential gene expression analysis between individual NK cell clusters in control and MM revealed no major transcriptomic differences. Rather, comparison of transcriptomic profiles of total BM NK cells from MM patients and controls revealed that NDMM patients had an NK cell profile with a reduction in cytotoxicity genes and an enrichment for genes involved in responses to cytokines and type I interferon signaling pathways. Our study suggests that alterations in NK cell subset composition is another likely mechanism of NK cell dysfunctionality in MM. This is supported by our findings that the altered NK cell repertoire is accompanied by impaired responses to therapeutic antibodies ex vivo. While several recent studies also seem to suggest NK cell subset variation^38, 39, 40^, further investigations are warranted into the mechanisms underlying the altered NK cell composition in MM and the time of onset of changes in the NK cell landscape, especially in MGUS and SMM.

Accumulating evidence in other types of cancer suggests that balanced presence of CD56^dim^ and CD56^bright^ NK cell subsets is biologically important and alteration of this ratio in peripheral blood^41,42^ and BM^38^ are associated with clinical outcome of MM patients, though these studies were small. In our study we did not focus on associations with clinical outcome, as currently our sample size is insufficiently large to establish clinical associations. Here, we focused on the effects of the relative decrease in abundance of cytotoxic NK cells, leading to reduced *ex vivo* responses to therapeutic antibodies in up to 20% of patients. Nonetheless, the relative increase in CD56^bright^ NK cell is likely to have importance as well. CD56^bright^ NK cells are important in shaping tumor microenvironments, as they impact function and recruitment of other immune subsets such as dendritic cells, T cells, macrophages, neutrophils, and endothelial cells into the tumor microenvironment through the production of pro-inflammatory cytokines and chemokines ^43^. Additionally, CD56^bright^ NK cells demonstrate tumor-associated pro-angiogenic activities^43^. However, the mechanisms underlying effects of CD56^bright^ NK cells on individual tumor microenvironment components remain poorly described, and while it is likely that the relative increase in CD56^bright^ NK cells impacts the MM BM microenvironment, how the CD56^bright^-microenvironment axis is implicated in MM pathobiology remains to be determined.

In relapsed patients, persistent loss of cytotoxic NK cells was accompanied by an expansion of IFN-responsive NK cells and increased activation of both CD56^bright^ and CD56^dim^ compartments. The increase in IFN-responsive NK cells is consistent with previous studies showing expansion of other IFN-responsive immune subsets in MM bone marrow^22, 44, 45^. Yet, the physiological function of these IFN-responsive NK cells, and their contribution to MM pathobiology, remains to be elucidated. In general, data on NK cells from relapsed/refractory MM (RRMM) patients is limited, yet previously reported gene expression profiles indicated increased activation and reduced negative regulation of NK cell activation, suggesting more chronic NK cell activation in the BM of RRMM patients. In functional assays, PB NK cells from RRMM are relatively unresponsive to elotuzumab^46^, which is in line with the inferior responses to therapeutic antibodies *in vitro* we describe for NDMM patients with persistent cytotoxic NK cell reduction. In a study of RRMM patients receiving daratumumab monotherapy, reduced frequencies of cytotoxic GZMB^+^CD56^dim^ NK cells were found in patients with primary and acquired daratumumab resistance and associated with inferior PFS and OS^18^, further highlighting that alterations in BM NK cell subset composition can contribute to difference in response to therapeutic antibodies in MM.

In summary, our data highlight the dynamic alterations of the BM NK cell compartment, including a persistent loss of cytotoxic BM NK cells in MM patients at diagnosis, during treatment and at disease relapse. Whether BM NK cell composition affects clinical outcomes and therapy responses, specifically in the context of daratumumab-containing therapy, warrants further study in a larger cohort. NK cell profiling could hold the potential of informing on clinical impact of immune-based therapies, which would be especially relevant in those therapeutic interventions where ADCC is the major mode-of-action.

## Methods

Cryopreserved bone marrow aspirates from 6 NDMM, 21 post-therapy and 6 RRMM patients were thawed and sorted into CD45^+^CD38^−^CD34^−^CD235a^−^CD71^−^ and CD45^+^CD38^+^CD34^−^CD235a^−^CD71^−^ immune fractions and processed for single-cell RNA sequencing according to the 10X Genomics protocol as previously described^22^. This dataset was supplemented with data from 5 controls and 13 additional NDMM patients that were previously generated in our lab^22^, resulting in a total single-cell dataset of 51 individuals, including 19 NDMM patients and 6 RRMM patients. As the majority of NK cells are CD38^+^^47, 48^ (Supplemental Figure 1A-B, Supplemental Figure 6A-B) and CD38^high^ NK cells have increased cytotoxic function against cancer cells^37^, CD38^+^ NK cells were identified *in silico* by transcription of *KLRF1, KLRD1, GNLY* and *NKG7* (Supplemental Figure 1) as previously described^26^. The single-cell NK cell dataset is available for interactive exploration at https://www.bmbrowser.org/. Demographics and clinical characteristics of MM patients and controls are described in Supplemental Table 1 and the study workflow is shown in Figure 1A. Additional experimental details are included in Supplemental Methods. All statistical analyses were performed using GraphPad Prism 7 software. Mann-Whitney U test was used to compare groups, and significance was set at P≤0.05. All graphs show median unless otherwise indicated in the figure legends.

## Acknowledgments

We thank members of Myeloma Research Rotterdam and the Department of Hematology for helpful discussions and input on the manuscript. This work was supported by grants from the European Union’s Horizon 2020 research and innovation program under the Marie Sklodowska-Curie grant agreement no. 70704 (to Z.K.).

## Author Contributions

S.T., T.C. and P.S. conceptualized the study; S.T., M.D.J., C.F. and T.C. developed the methodology; S.T., M.D.J., C.F., N.P. and E.S conducted the investigations. S.T., M.D.J., C.F., N.P., Z.K., G.v.B., R.H., K.N., E.S., M.D. analyzed the data; M.V. and M.v.D. collected the blood samples and clinical information, M.v.D., P.W., M.D., F.G., V.H.J.v.d.V., M.S., S.Z., N.D. and A.B. provided resources for the study, S.T. wrote the original draft, and all authors reviewed and edited the final manuscript. T.C and P.S. acquired funding and supervised the study.

## Conflict-of-interest Disclosures

M.D. is on the advisory board of Sanofi and GSK. F.G. received honoraria from and is on the advisory board of Janssen, Takeda, GSK, BMS/Celgene, Oncopeptides, Sanofi, Amgen, Pfizer, Roche and Abbvie. V.H.J.v.d.V. is an inventor on the EuroFlow-owned patent PCT/NL2013/0505420 (Methods, reagents, and kits for detecting minimal residual disease). The related patent is licensed to Cytognos (Salamanca, ES), who pays royalties to the EuroFlow Consortium. V.H.J.v.d.V. reports Laboratory Services Agreements with Agilent Technologies, Navigate, Cytognos, Janssen and BD Biosciences. S.Z. receives research support from Janssen and Takeda and is on the advisory boards for Janssen, BMS, Takeda, Oncopeptides and Sanofi without receiving personal fees. N.W.C.J.v.d.D. has received research support from Janssen Pharmaceuticals, AMGEN, Celgene, Novartis, Cellectis and BMS, and serves in advisory boards for Janssen Pharmaceuticals, AMGEN, Celgene, BMS, Takeda, Roche, Novartis, Bayer, Adaptive, and Servier, all paid to institution. A.B. received honoraria and/or is on the advisory board of Amgen, Sanofi, BMS and Janssen. P.S. consults for and receives research support from BMS/Celgene and Janssen.

## Notes

https://www.bmbrowser.org/

