## Supplemental Appendix for "Single-cell transcriptomic analysis of NK cell dynamics in myeloma patients reveal persistent reduction of cytotoxic NK cells from diagnosis to relapse"

Tahri et al.

**SUPPLEMENTARY APPENDIX**

**Table of Contents**


| Section | | Page |
|---|---|---|
| Methods | Supplementary Methods | 3 |
| Table S1 | Clinical characteristics patients single-cell RNA sequencing cohort | 8 |
| Table S2 | Differentially expressed genes (DEGs) between NK cell clusters of NDMM patients vs non-cancer controls | 10 |
| Table S3 | Signatures identified by Non-Matrix Factorization | 12 |
| Table S4 | Differentially expressed genes (DEGs) between NK cell clusters of NDMM patients with high vs low cytotoxicity | 13 |
| Figure S1 | Annotation of the $CD38^+$ NK cell dataset by single-cell RNA sequencing | 17 |
| Figure S2 | Cohort overview illustrating discovery cohort (single-cell RNA sequencing) and validation cohort (flowcytometry) analysed for *GZMK*$^+$CD56$^{bright}$ and *GZMB*$^+$CD56$^{dim}$ levels in the BM of NDMM patients | 19 |
| Figure S3 | Signature analysis by Non-Matrix Factorization | 19 |
| Figure S4 | Subset of NDMM patients have reduced frequencies of cytotoxic NK cells | 21 |
| Figure S5 | Dynamic alterations of the NK cell compartment during antitumor therapy and at relapse | 22 |
| Figure S6 | $CD38^+$ expression on NK cells in the BM of NDMM patients | 23 |
| Figure S7 | Annotation of the $CD38^-$ NK cell dataset by single-cell RNA sequencing | 24 |
| Figure S8 | Graphical abstract | 25 |

### SUPPLEMENTARY METHODS

#### Patient samples and cohorts

BM aspirates from patients with newly diagnosed MM were collected at the Erasmus Medical Centre (Rotterdam, the Netherlands) as part of inclusion into Dutch-Belgian Cooperative Trial Group for Hematology Oncology (HOVON) trials or European Myeloma Network (EMN) trials. The protocols were approved by the Institutional Review Board of the Erasmus Medical Centre (Rotterdam, The Netherlands). Patient samples were obtained after informed consent, in accordance with the Declaration of Helsinki. Control BM was obtained by sternal aspiration from donors undergoing cardiothoracic surgery or by manual BM collection from femur heads collected after hip replacement surgery. Bone marrow mononuclear cells (BMMCs) were isolated by density gradient centrifugation using Histopaque (Sigma-Aldrich) and subsequent plasma cell isolation was performed using magnetic $CD138^+$ bead enrichment (Stemcell). Plasma cells, pre-isolation fractions and plasma-cell depleted fractions were viably cryopreserved in 10% dimethyl sulfoxide (DMSO, Sigma-Aldrich).

BM samples for the single-cell transcriptomics cohort were obtained from patients included in the randomized phase 3 Cassiopeia trial which investigates the impact of daratumumab in combination with bortezomib, thalidomide and dexamethasone as first line treatment for transplant-eligible NDMM patients (HOVON-131/ IFM 2015-01; NCT02541383)[1, 2]. BM samples for flow cytometric analysis of NK cell subsets were obtained from patients included in a study of an anti-CD38 monoclonal antibody in transplant-eligible NDMM patients or in the phase II trial of Ixazomib-Daratumumab-low-dose-Dexamethason (Ixa-Dara-Dex) in frail NDMM patients (HOVON-143; NTR6297)[3].

#### Single-cell transcriptomics

Cryopreserved BM aspirates were thawed by serial dilution in complete media (DMEM + 10%FCS) with 10 mg/ml DNAse (Sigma-Aldrich) according to the 10X Genomics protocol. Fc receptors were blocked with 10% normal human AB serum (Sigma-Aldrich). Cells were labelled for 20-30 min on ice with FACS staining buffer (PBS containing 2% FCS) with a mixture of the following antibodies: CD235a-PECy7 (1:20; HI264, BioLegend), CD34-PECF610 (1:100; 4H11, eBioscience), CD45-APC

(1:20; 2D1, eBioscience), CD38-FITC (1:20; MHCD3801, Life Technologies), CD71-Alexa Fluor 700 (1:20; MEM-75, ExBio) and 7AAD (1:100; Beckman Coulter) or DAPI (1:100; Life Technologies). $CD45^{+}CD38^{-}CD34^{-}CD235a^{-}CD71^{-}$ and $CD45^{+}CD38^{+}CD34{-}CD235a^{-}CD71^{-}$ immune fractions were sorted using a FACSAria III (BD Biosciences) and immediately processed for single-cell transcriptomics. The 10X Chromium Single cell 3’ reagent Kit v3 (10X Genomics) was used for construction of single-cell transcriptomic libraries. Reverse transcription, cDNA amplification, and library preparation were all performed according to the manufacturer’s instructions. The final library construct was assessed by the Agilent Bioanalyzer High Sensitivity Chip (Agilent) for quality control. Libraries were sequenced using a NovaSeq6000 platform (Illumina) at a sequencing depth of ~ 20.000 reads per cell.

**Single-cell transcriptomics analysis**

Raw fastq files were processed with 10X Genomics CellRanger (version 3.0.2., 10X Genomics), which included demultiplexing, alignment, filtering, barcode and unique molecular identifier (UMI) counting. Subsequent analysis was performed using the Seurat 4.2.0 package[4] in R 4.2.2. We performed ambient RNA correction with SoupX[5]. We first processed each individual patient separately prior to combining data from multiple patients. Low-quality cells were removed when expressing less than 200 genes, mitochondrial genes comprised more than 10% of the total transcript pool or high feature counts were observed (doublets). Dimensionality reduction was achieved for each dataset by selecting the 2,000 most variable genes and performing principal component analysis (PCA). Cells were clustered using the Louvain algorithm in Seurat’s FindClusters function. Finally, cells were visualized in a UMAP embedding. Single cell transcriptomic data from individual patients were integrated, based on the identified anchors using the IntegrateData function in Seurat. Afterwards we repeated scaling, normalization, PCA, clustering and UMAP embedding for the integrated dataset. NK cells were identified based on transcription of *KLRF1*, *KLRD1*, *NKG7* and *GNLY*[6]; these cells were extracted, thereby obtaining 24,664 BM NK cells for further analysis. We assessed well-defined marker genes across clusters to identify potential contaminating cell populations such as T cells (*CD8A, CD8B, CD4*), B cells (*MS4A1, CD19, VPREB1*), plasma cells (*SDC1, LAMP5, SLAMF7*) and myeloid cells (*LYZ, CD14,*

*FCGR3B, ELANE, FCER1A, CD1C*). One cluster containing a cell type admixture was removed. Furthermore, immunoglobulin and hemoglobin genes were removed prior to subsequent analysis. Finally, we performed differential expression analysis on the integrated dataset to identify genes significantly upregulated in each cluster compared to all other clusters (FDR-adjusted $p \leq 0.05$) using the Seurat function FindMarkers. Pseudotime analysis was carried out using Monocle 3[7]. Cytotoxicity and cytokine-producing scores were calculated based on selected genes using the AddModuleScore function of Seurat.

**Analysis of CD38$^-$ dataset**

The CD38$^-$ dataset contained 9,354 NK cells distributed among 3 transcriptional clusters, including a cytotoxic GZMB$^+$CD56$^{dim}$ NK cell cluster and a naïve and activated (ltNK-like) GZMK$^+$CD56$^{bright}$ NK cell cluster (Supplemental Figure 7A-C), similarly to the CD38$^+$ NK cell dataset. Due to the limited number of cells per patient (<180 cells/ patient) we refrained from further analyzing this dataset.

**Signature analysis by Non-Negative Matrix Factorization**

Signature analysis was performed using the regularized non-negative matrix factorization algorithm[8]. For a reduction in compute time a GPU specific implementation was used (https://github.com/broadinstitute/SignatureAnalyzer-GPU). The regularized non-negative matrix factorization algorithm requires a count matrix based on the most variable genes as the input. Variable genes represented the rows and single cells represented the rows of the count matrix. The optimization algorithm was allowed to run for 10000 iterations at maximum. The default exponential prior was used for both factorization components (i.e., gene signatures and activity scores). Seurat was used for creating UMAP plots based on the final factorization.

**Flow cytometry**

Fresh or cryopreserved BMMCs were stained using FACS staining buffer (PBS; Gibco™ + 2% FCS; Corning). Fc receptors were blocked with human TruStain FcX (422301, Biolegend) according to the manufacturers protocol. Cells were stained with the indicated fluorochrome-conjugated antibodies for 20-30 min on ice. For intracellular staining, surface-stained cells were fixed and permeabilized using Cyto-Fast Fix/Perm Buffer set (Biolegend) according to the manufacturer's instructions. Multicolor flow cytometry was performed using an LSR II flow cytometer (BD Biosciences) or BD Symphony Flow cytometer and analyzed by FlowJo V10 Software. The gating strategies are summarized in Supplementary Figure S3 and Figure S7.

**ADCC and functional assay**

Cryopreserved bone marrow nuclear cells (BMMCs) from untreated NDMM patient samples were thawed by serial dilution in complete media (DMEM + 10%FCS) with 10 mg/ml DNAse. NK cells were isolated by negative selection using the EasySep Human NK cell Isolation Kit (#17955, STEMCELL). Enriched NK cells were checked for purity and then cultured for 12-24 hours in RPMI-1640 containing 2mM L-glutamine and 100 ug/ml penicillin and streptomycin (Gibco) supplemented with 10% fetal calf serum (FCS; Corning), and 200-300 IU/ml of recombinant human IL-2 (rhIL-2) (PeproTech) at a density of $2x10^6$/mL. The rhIL-2 stimulated NK cells were washed with PBS (Gibco™) before they were co-cultured with target cells.

MM.1S, RPMI-8226 myeloma cell lines and K562 (ATCC) were used as target cells in the co-culture assays with NK cells and cultured in RPMI1640 (Gibco) supplemented with 10% FCS and 1% penicillin/streptomycin (Gibco). All cell lines were mycoplasma free (MycoAlert Detection Kit) and cells were cultured at 37C and 5% CO2. $2 \times 10^5$ cells were added to each well of a U-bottom 96-well plate and pre-incubated for 30 min with 1 ug/ml or 10 ug/ml daratumumab (Creative Biolabs) or human IgG (BioXcell). Target cells were washed twice with FACS staining buffer and then co-cultured with NK cells in various effector to target ratios (1:2, 1;1, 2:1) for 4 hours in the presence of CD107a-APC (clone H4A3, Biolegend). Cells were then harvested, resuspended in FACS buffer (PBS; Gibco™ + 2% FCS; Corning) and DAPI (1:500, Life Technologies) and NK cell markers were added (CD3-AF700, clone UCHT1, Biolegend, CD56-PE clone 5.1H11 (Biolegend), CD16-FITC clone CD2F1 (BD Pharmingen)).

**Statistical analysis**

All statistical analyses were performed using GraphPad Prism 7 software. Mann-Whitney U test was used to compare groups, and a P≤0.05 was considered statistically significant. All figures are representative of (or pooled from) at least two in vitro experiments. All graphs show median unless otherwise indicated in the figure legends.

**Data availability**

The single cell RNA-sequencing dataset described in this study are available on ArrayExpress no. E-MTAB-13157. The single-cell RNA sequencing dataset of NK cells can be interactively explored at https://www.bmbrowser.org/. The scripts generated during this study will be available through GitHub at https://github.com/MyelomaRotterdam/Tahri-et-al.-2023.

### SUPPLEMENTARY TABLES

**Supplemental Table 1. Clinical characteristics patients single-cell RNA sequencing cohort**

| Sample ID | Timepoint | Gender | Age | Cytogenetics | R-ISS | Plasma cells in BM (%) | M-protein | Serum M-protein (g/dl) |
|---|---|---|---|---|---|---|---|---|
| CBM1 | N/A | F | 80 | N/A | N/A | N/A | N/A | N/A |
| CBM2 | N/A | M | 79 | N/A | N/A | N/A | N/A | N/A |
| CBM3 | N/A | M | 76 | N/A | N/A | N/A | N/A | N/A |
| CBM4 | N/A | F | 76 | N/A | N/A | N/A | N/A | N/A |
| CBM5 | N/A | M | 64 | N/A | N/A | N/A | N/A | N/A |
| MM1 | Newly diagnosed | F | 58 | HD | III | 18.9 | IgG | 0.35 |
| MM2 | Newly diagnosed | M | 61 | HD | II | 11.5 | IgG | 3.72 |
| MM3 | Newly diagnosed | M | 63 | HD | II | 13.8 | IgG | 2.58 |
| MM4 | Newly diagnosed | M | 59 | HD | I | 3.19 | IgG | 2.13 |
| MM5 | Newly diagnosed | M | 64 | HD | I | 1.2 | IgG | 2.84 |
| MM6 | Newly diagnosed | F | 63 | HD | I | 0.9 | IgG | 2.04 |
| MM7 | Newly diagnosed | M | 58 | t(4;14) | II | 20.1 | Kappa | 0.99 |
| MM8 | Newly diagnosed | M | 61 | t(4;14) | II | 4.6 | IgA | 3.09 |
| MM9 | Newly diagnosed | F | 61 | t(4;14) | II | 5.5 | IgG | 1.28 |
| MM10 | Newly diagnosed | F | 64 | t(11;14), del17p | III | 6 | IgG | 3.85 |
| MM11 | Newly diagnosed | M | 64 | t(11;14) | I | 16.3 | IgG | 2.23 |
| MM12 | Newly diagnosed | F | 60 | t(11;14) | II | 18.4 | Kappa | 0.41 |
| MM13 | Newly diagnosed | F | 54 | del17p | II | 9.2 | IgG | 0.22 |
| MM14 | Newly diagnosed | M | 36 | HD, 1q amplification | III | N/A | IgG | 6.38 |
| MM15 | Newly diagnosed | F | 58 | HD, t(14;20) | III | N/A | IgA | 2.93 |

| | | | | | | | | |
|---|---|---|---|---|---|---|---|---|
| MM16 | Newly diagnosed | M | 52 | N/A | II (ISS) | N/A | IgA | 1.1 |
| MM17 | Newly diagnosed | F | 60 | N/A | III (ISS) | N/A | IgG | 5.5 |
| MM18 | Newly diagnosed | F | 47 | N/A | I (ISS) | N/A | IgG | 3.4 |
| MM19 | Newly diagnosed | F | 62 | del17p | III | N/A | IgG | 6.6 |
| MM20 | C4D28 | M | 64 | t(11;14) | II | 4.08¥ | N/A | N/A |
| MM21 | C4D28 | F | 65 | Hyperdiploid | II | 0.51¥ | IGG | 5.6 |
| MM22 | C4D28 | M | 59 | Hyperdiploid | I | 0.002¥ | N/A | N/A |
| MM23 | D100 | F | 65 | Hyperdiploid | II | 0.0016¥ | IGG | 7.88 |
| MM24 | D100 | M | 60 | Normal* | I | 0¥ | N/A | N/A |
| MM25 | D100 | M | 47 | Hyperdiploid | II | 0.0038¥ | IGG | 4.11 |
| MM26 | D100 | F | 52 | Hyperdiploid | II | 0¥ | IGG | 3.59 |
| MM27 | D100 | M | 59 | Hyperdiploid | I | 0¥ | IGG | 3.48 |
| MM28 | D100 | M | 64 | Hyperdiploid | I | 0¥ | IGG | 6.47 |
| MM29 | W25 | M | 64 | Hyperdiploid | I | 0.002¥ | IGG | N/A |
| MM30 | W25 | M | 62 | t(11;14) | I | N/A | N/A | N/A |
| MM31 | W52 | F | 57 | Hyperdiploid | I | 0¥ | IGG | N/A |
| MM32 | W52 | M | 55 | Hyperdiploid | II | 0¥ | IGA | N/A |
| MM33 | W52 | F | 62 | t(11;14) | I | 0.49¥ | N/A | N/A |
| MM34 | W52 | F | 55 | t(11;14) | I | 0.003¥ | N/A | N/A |
| MM35 | W52 | M | 63 | Hyperdiploid | II | 0.71¥ | IGG | N/A |
| MM36 | W105 | M | 40 | t(4;14) | II | 0.007¥ | IGG | N/A |
| MM37 | W105 | F | 52 | Hyperdiploid | II | 0¥ | IGG | N/A |
| MM38 | W105 | M | 55 | t(11;14) | II | 0¥ | N/A | N/A |
| MM39 | W105 | M | 63 | Hyperdiploid | II | 8.75¥ | IGG | N/A |
| MM40 | W105 | M | 55 | Hyperdiploid | II | 0¥ | IGA | N/A |
| MM41 | Relapse | F | 56 | Hyperdiploid | II | N/A | IGG | N/A |
| MM42 | Relapse | M | 40 | t(4;14) | II | 0.2¥ | IGG | N/A |
| MM43 | Relapse | F | 53 | t(4;14) | II | 10.2¥ | IGG | N/A |
| MM44 | Relapse | M | 43 | t(4;14) | II | 0.1¥ | IGA | N/A |
| MM45 | Relapse | M | 45 | Hyperdiploid | II | 0.3¥ | IGG | N/A |
| MM46 | Relapse | F | 65 | Hyperdiploid | II | 0.1¥ | IGG | N/A |

* Del17p not evaluable. ¥ Determined by EuroFlow MRD flowcytometry.

**Supplemental Table 2. Differentially expressed genes (DEGs) between NK cell clusters of NDMM patients vs non-cancer controls.** Positive Log2FoldChanges denote upregulated genes in non-cancer controls, negative log2FoldChanges denote downregulated genes in non-cancer controls (and thus upregulated in NDMM patients). Only genes with an adjusted p-values ≤ 0.05 are presented.

| **Cluster** | **Gene name** | **Log2Fold Change** | **Adjusted p-value** |
|---|---|---|---|
| Conventional cytotoxic $CD56^{dim}$ NK cells | IL32 | 0.341 | 3.11E-04 |
| Transitional/active NK cells | PYHIN1 | 0.547 | 3.88E-04 |
| | CMC1 | 0.420 | 2.60E-03 |
| | NFKBIA | 0.387 | 1.75E-02 |
| | AGTRAP | 0.370 | 1.60E-03 |
| | GZMM | 0.283 | 1.88E-03 |
| | LINC00152 | 0.240 | 1.64E-02 |
| | ACTG1 | 0.232 | 3.21E-03 |
| | ZYX | 0.226 | 8.58E-03 |
| | MT2A | -0.297 | 1.22E-02 |
| Activated $CD56^{bright}$ | ZFP36 | 0.547 | 3.48E-02 |
| | NEAT1 | 0.505 | 1.73E-02 |
| | HIST1H1E | 0.362 | 1.71E-03 |
| | GPX1 | 0.352 | 4.18E-02 |
| | PRDX2 | 0.344 | 3.41E-04 |
| | MT1X | 0.318 | 5.35E-04 |
| | AREG | 0.293 | 2.00E-03 |
| | FGFBP2 | 0.220 | 7.97E-03 |
| | GZMM | 0.215 | 3.39E-02 |
| | TTN | 0.206 | 2.31E-04 |
| | LYZ | -0.223 | 6.17E-03 |
| IFN-responsive NK cells | CLIC3 | 0.599 | 4.85E-04 |
| | PLEC | 0.487 | 5.16E-04 |
| | PLAC8 | 0.476 | 1.59E-03 |
| | NEAT1 | 0.428 | 1.47E-03 |
| | PYCARD | 0.380 | 1.46E-04 |
| | HOPX | 0.377 | 6.65E-04 |
| | FOSB | 0.352 | 3.10E-02 |
| | CCL4 | 0.326 | 4.59E-02 |
| | KLRD1 | 0.281 | 9.00E-04 |
| | FGR | 0.243 | 7.93E-03 |
| | ANXA5 | -0.203 | 2.46E-04 |
| | EZH2 | -0.212 | 1.08E-03 |
| | SOCS3 | -0.233 | 1.54E-04 |

| | | | |
|---|---|---|---|
| | HIST1H4C | -0.257 | 3.25E-02 |
| | GADD45A | -0.262 | 1.59E-04 |
| | SAMSN1 | -0.273 | 1.31E-04 |
| | TNFAIP3 | -0.286 | 1.91E-02 |
| Naive CD56$^{bright}$ | IL32 | 0.597 | 3.74E-03 |
| | SELL | 0.575 | 7.71E-03 |
| | JUN | 0.520 | 3.62E-02 |
| | FCGR3A | 0.513 | 3.02E-02 |
| | NEAT1 | 0.427 | 4.65E-03 |
| | MT2A | 0.423 | 4.88E-02 |
| | HLA-DPB1 | 0.405 | 1.03E-03 |
| | TRBC1 | 0.397 | 1.85E-02 |
| | SAMHD1 | 0.395 | 5.01E-04 |
| | GRASP | 0.371 | 4.44E-04 |
| | C1orf162 | 0.339 | 1.22E-02 |
| | AREG | 0.338 | 2.97E-02 |
| | HIST1H1E | 0.283 | 9.11E-03 |
| | TRGC2 | 0.274 | 3.54E-02 |
| | IER2 | 0.270 | 2.83E-02 |
| | SYNE1 | 0.262 | 8.95E-03 |
| | TUBB4B | 0.251 | 1.03E-02 |
| | GAPDH | 0.229 | 8.32E-03 |
| | LMNB1 | 0.224 | 6.06E-03 |
| | XCL2 | -0.275 | 5.12E-03 |
| Proliferating NK cells | CENPA | 0.357 | 5.72E-03 |
| | PTGER4 | 0.338 | 4.25E-04 |
| | SGOL1 | 0.240 | 4.91E-02 |
| | PKMYT1 | 0.223 | 1.10E-03 |
| | CENPE | 0.207 | 2.57E-03 |
| | IGFBP7 | -0.280 | 3.50E-03 |
| Terminal NK cells | NFKBIZ | -0.218 | 5.50E-03 |
| | HN1 | -0.225 | 2.30E-03 |
| | CDC42EP3 | -0.237 | 3.87E-04 |
| | HIST1H2BF | -0.298 | 5.17E-03 |
| | DUT | -0.470 | 8.80E-03 |

**Supplemental Table 3. 15 gene signatures (factors) identified by Non-Matrix Factorization.**

| Signature | Biological interpretation | Top genes |
|---|---|---|
| Signature 1 | MM contamination | *IGLC2, IGLC3, JCHAIN, IGHM* |
| Signature 2 | Cytotoxicity/*CCL5* high | *CCL5, NKG7, S100A4* |
| Signature 3 | Activation | *DUSP2, ZFP36, MAP3K8* |
| Signature 4 | IFN-responsive | *PMAIP1, ISG15, IFIT2, TNFAIP3, IFIT3* |
| Signature 5 | Activation/*CCL3* high | *CCL3, CCL4, CCL3L3, CD69* |
| Signature 6 | Activation/immediate early genes | *FOS, JUN, NFKBIA, IER2* |
| Signature 7 | Activated $CD56^{bright}$ | *PTMA, CD7, NFKBIA, NKG7, TYROBP, ACTG1, CSTW, GSTP1, XCL2* |
| Signature 8 | MM contamination | *IGKC, JCHAIN, SSR4* |
| Signature 9 | $CD56^{dim}$ | *PTMA, NKG7, FTL, MT2A, FGFBP2, S100A6, HMGB1, S100A4* |
| Signature 10 | Cytotoxicity | *NKG7, GZMB, FGFBP2, CST7, PTMA, GZMA, CTSW, SPON2, S100A4, ACTG1, PRF1* |
| Signature 11 | *FTH1* high | *FTH1, PTMA, CD7* |
| Signature 12 | Unknown | *KLRB1, NKG7, CMC1, NEAT1, HMGB1, RUNX3, TMEM2, KLRD1, KLRF1* |
| Signature 13 | *F1TX* high/ contamination | *H1FX, MPO, H1F0* |
| Signature 14 | Proliferating | *PTMA, PIMA, GAPDH, TUBA1B, HMGN2, HMGB2, ACTG1, HMGB1, STMN1* |
| Signature 15 | *GNLY* high | GNLY, NKG7, CTSW |

**Supplemental Table 4. Differentially expressed genes (DEGs) between NK cell pool of NDMM patients with high vs low cytotoxicity.** Positive log2FoldChanges denote upregulated genes in NDMM patients with high cytotoxicity, negative log2FoldChanges denote downregulated genes in NDMM patients with high cytotoxicity (and thus upregulated in NDMM patients with low cytotoxicity). Only genes with an adjusted p-values ≤ 0.05 are presented.

| Gene name | Log2Fold Change | Adjusted p-value |
|---|---|---|
| IFIT3 | 0.619 | 1.43E-09 |
| PMAIP1 | 0.607 | 4.72E-10 |
| SYNE1 | 0.439 | 1.21E-09 |
| HIST1H4C | 0.416 | 1.62E-30 |
| CEP78 | 0.361 | 2.41E-20 |
| ARL4C | 0.349 | 8.64E-70 |
| HOPX | 0.348 | 2.88E-07 |
| AHNAK | 0.334 | 7.70E-17 |
| MT1X | 0.325 | 8.82E-158 |
| TOB1 | 0.320 | 8.31E-35 |
| PTGER4 | 0.310 | 2.14E-22 |
| ITGAM | 0.305 | 1.30E-52 |
| MAF | 0.299 | 1.04E-124 |
| HIST1H1D | 0.287 | 2.62E-39 |
| HLA-DPB1 | 0.286 | 5.77E-05 |
| LINC00936 | 0.276 | 2.37E-160 |
| GZMB | 0.270 | 4.33E-82 |
| ARID5B | 0.252 | 2.22E-43 |
| HES6 | 0.251 | 1.65E-19 |
| SOD2 | 0.249 | 3.18E-71 |
| RGS2 | 0.239 | 3.85E-140 |
| SPON2 | 0.232 | 4.41E-53 |
| SLC2A3 | 0.230 | 1.48E-95 |
| HES4 | 0.228 | 2.13E-11 |
| TIMP1 | 0.227 | 1.76E-45 |
| SELL | 0.224 | 3.59E-25 |
| RP11-160E2.6 | 0.222 | 1.02E-58 |
| LMNB1 | 0.219 | 4.65E-02 |
| CDC42EP3 | 0.218 | 1.21E-108 |
| CD2 | 0.217 | 1.36E-41 |

| | | |
|---|---|---|
| HERC5 | 0.215 | 4.69E-61 |
| APLP2 | 0.214 | 2.47E-53 |
| TMEM107 | 0.212 | 1.95E-97 |
| HIST1H2AC | 0.207 | 8.67E-06 |
| PTMS | 0.206 | 3.96E-122 |
| PSAP | 0.201 | 3.38E-97 |
| ATP2B1 | 0.199 | 1.64E-31 |
| IRF2BP2 | 0.197 | 5.89E-90 |
| NAMPT | 0.193 | 1.21E-41 |
| KPNA2 | 0.187 | 2.55E-30 |
| ZFP36L1 | 0.181 | 4.28E-46 |
| EIF4A3 | 0.178 | 4.21E-106 |
| VMP1 | 0.175 | 1.01E-64 |
| CDC25B | 0.168 | 6.51E-49 |
| PRF1 | 0.167 | 1.49E-31 |
| TMPO | 0.164 | 8.37E-40 |
| SOX4 | 0.163 | 2.36E-113 |
| LCK | 0.162 | 3.79E-15 |
| RGS10 | 0.162 | 3.56E-38 |
| GPR65 | 0.158 | 1.95E-38 |
| DUT | 0.158 | 7.58E-85 |
| MXD1 | 0.155 | 3.64E-175 |
| CES1 | 0.155 | 3.89E-07 |
| SAMHD1 | 0.154 | 1.72E-125 |
| CITED2 | 0.151 | 6.71E-77 |
| FAM101B | 0.141 | 1.18E-156 |
| MT1E | 0.141 | 1.97E-114 |
| ID2 | 0.140 | 1.33E-30 |
| EZH2 | 0.139 | 3.27E-161 |
| FGFBP2 | 0.137 | 1.59E-78 |
| GADD45A | 0.136 | 7.88E-64 |
| SOCS3 | 0.136 | 8.21E-91 |
| ATF3 | 0.135 | 4.76E-58 |
| MIS18BP1 | 0.134 | 4.45E-36 |
| HIST1H2BC | 0.132 | 2.61E-188 |
| DEK | 0.130 | 1.03E-13 |
| HNRNPAB | 0.129 | 1.44E-157 |
| LDLR | 0.128 | 4.40E-134 |
| CDKN1C | 0.127 | 3.66E-121 |
| TRGC2 | 0.124 | 4.50E-17 |
| MARCKSL1 | 0.121 | 1.63E-190 |

| | | |
|---|---|---|
| MIR22HG | 0.121 | 2.14E-67 |
| KIF20B | 0.118 | 3.88E-183 |
| TKT | 0.118 | 5.11E-65 |
| GZMH | 0.116 | 4.18E-22 |
| PYCARD | 0.113 | 1.05E-13 |
| C10orf54 | 0.113 | 2.26E-40 |
| NFKBIZ | 0.112 | 5.20E-67 |
| PHLDA1 | 0.111 | 1.11E-101 |
| HIST1H2AG | 0.109 | 4.05E-52 |
| CST3 | 0.105 | 3.10E-40 |
| CTSS | 0.104 | 5.96E-26 |
| GMNN | 0.102 | 6.50E-114 |
| HIST2H2AC | 0.101 | 6.60E-155 |
| TRGC1 | 0.101 | 1.85E-13 |
| SPTBN1 | 0.100 | 8.00E-34 |
| FOSB | -0.104 | 3.69E-06 |
| PTMA | -0.111 | 2.22E-03 |
| SLBP | -0.116 | 1.29E-02 |
| IL2RB | -0.119 | 5.11E-42 |
| TFDP1 | -0.121 | 8.34E-11 |
| RGCC | -0.122 | 7.55E-08 |
| IFITM3 | -0.124 | 4.06E-118 |
| TTN | -0.142 | 3.12E-50 |
| STMN1 | -0.151 | 1.23E-29 |
| GADD45G | -0.156 | 7.08E-174 |
| MATK | -0.156 | 3.65E-15 |
| HOXA5 | -0.166 | 1.14E-48 |
| TMIGD2 | -0.167 | 3.64E-89 |
| DDIT4 | -0.171 | 6.29E-13 |
| CD160 | -0.173 | 2.64E-101 |
| KLRB1 | -0.208 | 3.65E-07 |
| PPP1R14B | -0.217 | 1.13E-11 |
| ITM2C | -0.230 | 8.61E-173 |
| H1FX | -0.255 | 1.76E-16 |
| MAP3K8 | -0.266 | 2.29E-05 |
| PDE4B | -0.277 | 8.56E-17 |
| IER2 | -0.296 | 6.81E-03 |
| CD96 | -0.344 | 4.61E-15 |
| CCL5 | -0.354 | 3.93E-04 |
| CD7 | -0.358 | 2.64E-61 |
| CMC1 | -0.360 | 3.05E-57 |

| | | |
|---|---|---|
| SPRY1 | -0.366 | 5.19E-17 |
| AREG | -0.416 | 2.51E-21 |
| GSTP1 | -0.441 | 8.17E-41 |
| TYROBP | -0.453 | 1.52E-40 |
| NFKBIA | -0.454 | 3.91E-21 |

### SUPPLEMENTARY FIGURE

**Supplemental Figure S1. Annotation of the CD38⁺ NK cell dataset. (**A) Transcription of NK cell makers *KLRD1*, *KLRF1*, *NKG7* and *GNLY* in CD38⁺ myeloma bone marrow immune dataset. (B) Transcription of T cell markers *CD3D*, *CD4*, *CD8A* and *CD8B* in CD38⁺ myeloma bone marrow immune dataset. (C) Representative flow cytometry gating strategy for intracellular Granzyme K and Granzyme B levels within $CD56^{bright}$ and $CD56^{dim}$ NK cells from bone marrow mononuclear cells (BMMCs) from aspirates of NDMM patients. (D) DotPlot of selected marker genes characterizing different NK cell clusters. Color indicates average expression of the gene and dot size indicates percent of cells in which expression of the gene is detected. (E) Violin plots demonstrating the expression of maturation markers for each cluster. (F) Dotplot depicting transcription of cytotoxicity genes among conventional cytotoxic $CD56^{dim}$ and terminal NK cells. (G) Cytoskeleton remodeling and ribosome signature as defined by Liu et al., depicted across NK cell clusters. (H) *GZMK* and *GZMB* transcription among IFN-responsive NK cells (left) and proliferating NK cells (right), split by controls vs myeloma patients. (I) ILC gene signature as defined by Bjorklund et al., depicted across NK cell clusters.

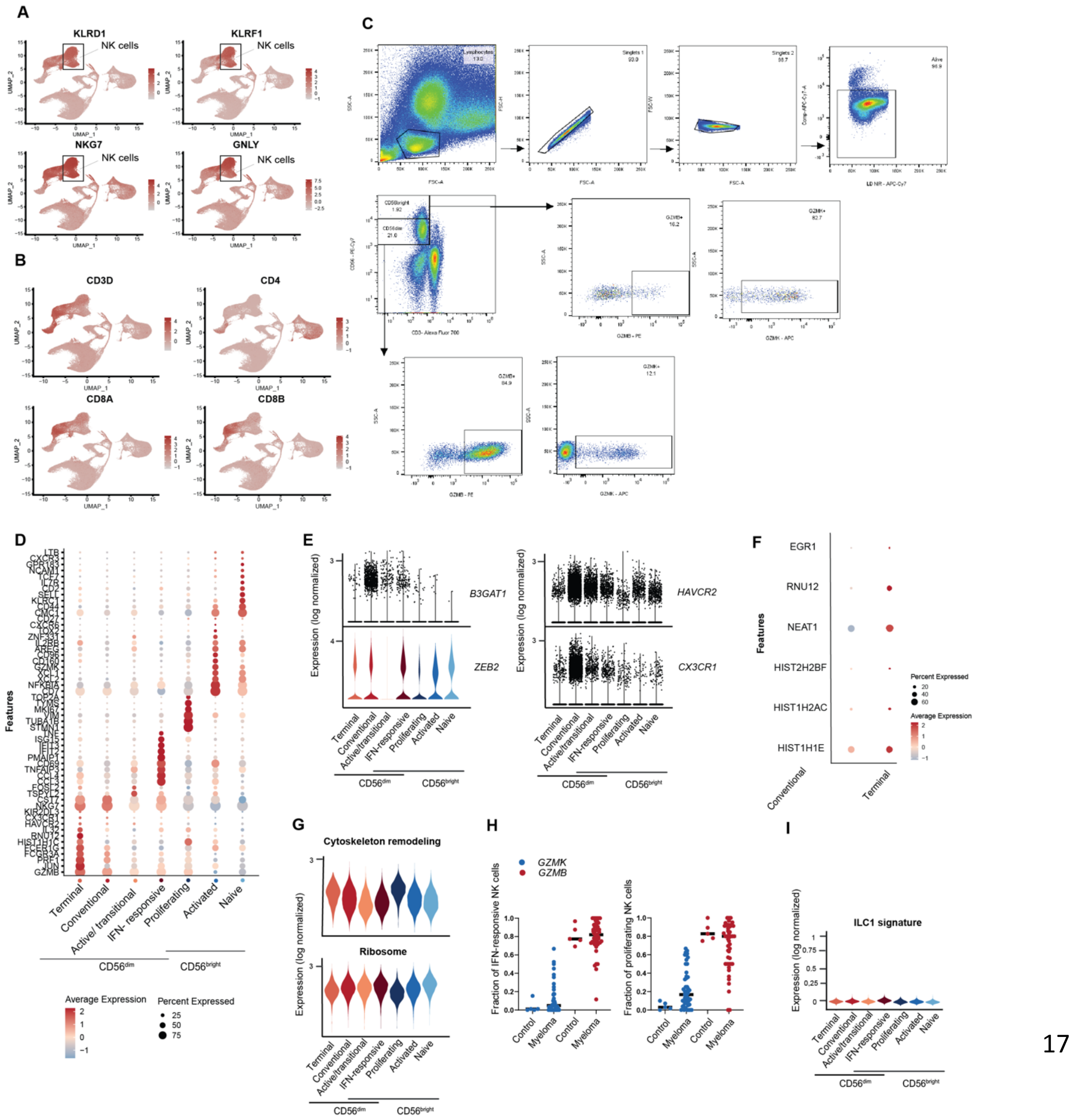

**Supplemental Figure S2. Cohort overview.** (A) Illustrating discovery cohort (single-cell RNA sequencing) and validation cohort (flowcytometry) analysed for *GZMK*$^{+}$CD56$^{bright}$ and *GZMB*$^{+}$CD56$^{dim}$ levels in the BM of NDMM patients. (B) Overview of treatment schedule per trial.

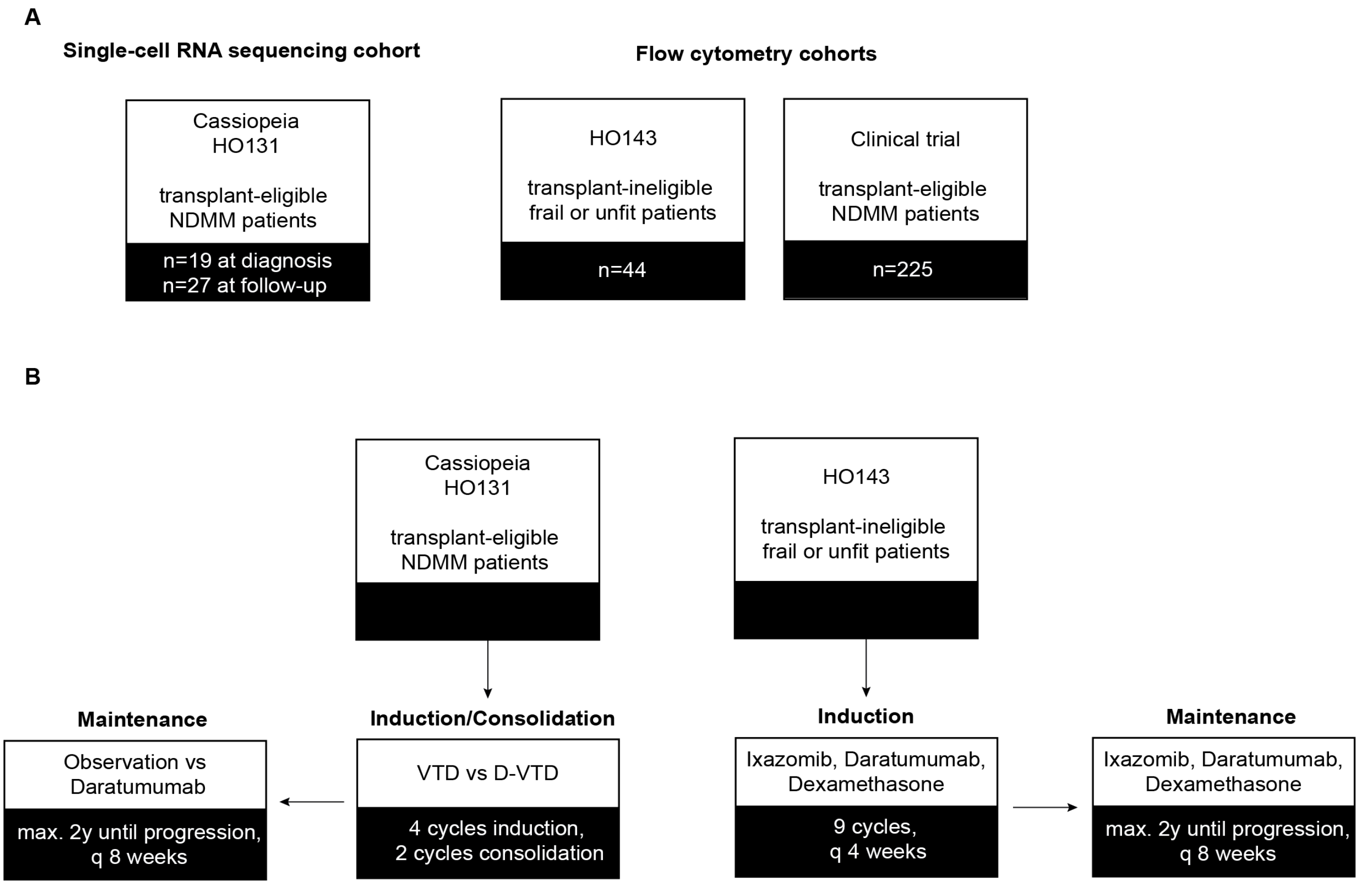

**Supplemental Figure S3. Non-Matrix-Factorization to identify NK cell gene signatures.** (A) Left: Transcription of gene signature 4 identifying IFN-responsive NK cells. Right: Violin plot depicting transcription of IFN-responsive gene signature across 7 NK cell clusters. (B) Left: Transcription of gene signature 7 activated $CD56^{bright}$ NK cells. Right: Violin plot depicting transcription of gene signature specific for activated $CD56^{bright}$ NK cells across 7 NK cell clusters. (C) Left: Transcription of gene signature 14 identifying proliferating NK cells. Right: Violin plot depicting transcription of proliferating gene signature across 7 NK cell clusters. (D) Upper: Bar plot depicting genes contributing to gene signature 3 encompassing activation genes. Lower left: Violin plot depicting transcription of gene signature 3 among control and NDMM NK cell clusters. Lower right: Violin plot depicting transcription of gene signature 3 across 7 NK cell clusters. (E) Upper: Bar plot depicting genes contributing to gene signature 6 encompassing activation genes. Lower left: Violin plot depicting transcription of gene signature 6 among control and NDMM NK cell clusters. Lower right: Violin plot depicting transcription of gene signature 6 across 7 NK cell clusters. (F) Upper: Bar plot depicting genes contributing to gene signature 10 encompassing cytotoxicity genes. Lower left: Violin plot depicting transcription of gene signature 10 among control and NDMM NK cell clusters. Lower right: Violin plot depicting transcription of gene signature 10 across 7 NK cell clusters. (G) Left: Violin plot depicting transcription of gene signature 14 encompassing proliferation gene signature across 7 NK cell clusters. Right: Violin plot depicting transcription of gene signature 14 across NDMM patients with high or low cytotoxicity. (H) Left: Violin plot depicting transcription of gene signature 9 encompassing $CD56^{dim}$ gene signature across 7 NK cell clusters. Right: Violin plot depicting transcription of gene signature 9 across NDMM patients with high or low cytotoxicity. (I) Left: Violin plot depicting transcription of gene signature 10 encompassing cytotoxicity gene signature across 7 NK cell clusters. Right: Violin plot depicting transcription of gene signature 10 across NDMM patients with high or low cytotoxicity. (J) Left: Violin plot depicting transcription of gene signature 15 encompassing *GNLY* high gene signature across 7 NK cell clusters. Right: Violin plot depicting transcription of gene signature 15 across NDMM patients with high or low cytotoxicity. (K) Left: Violin plot depicting transcription of gene signature 2 encompassing *CCL5* high gene signature across 7 NK cell clusters. Right: Violin plot depicting transcription of gene signature 2 across NDMM patients with high or low cytotoxicity.

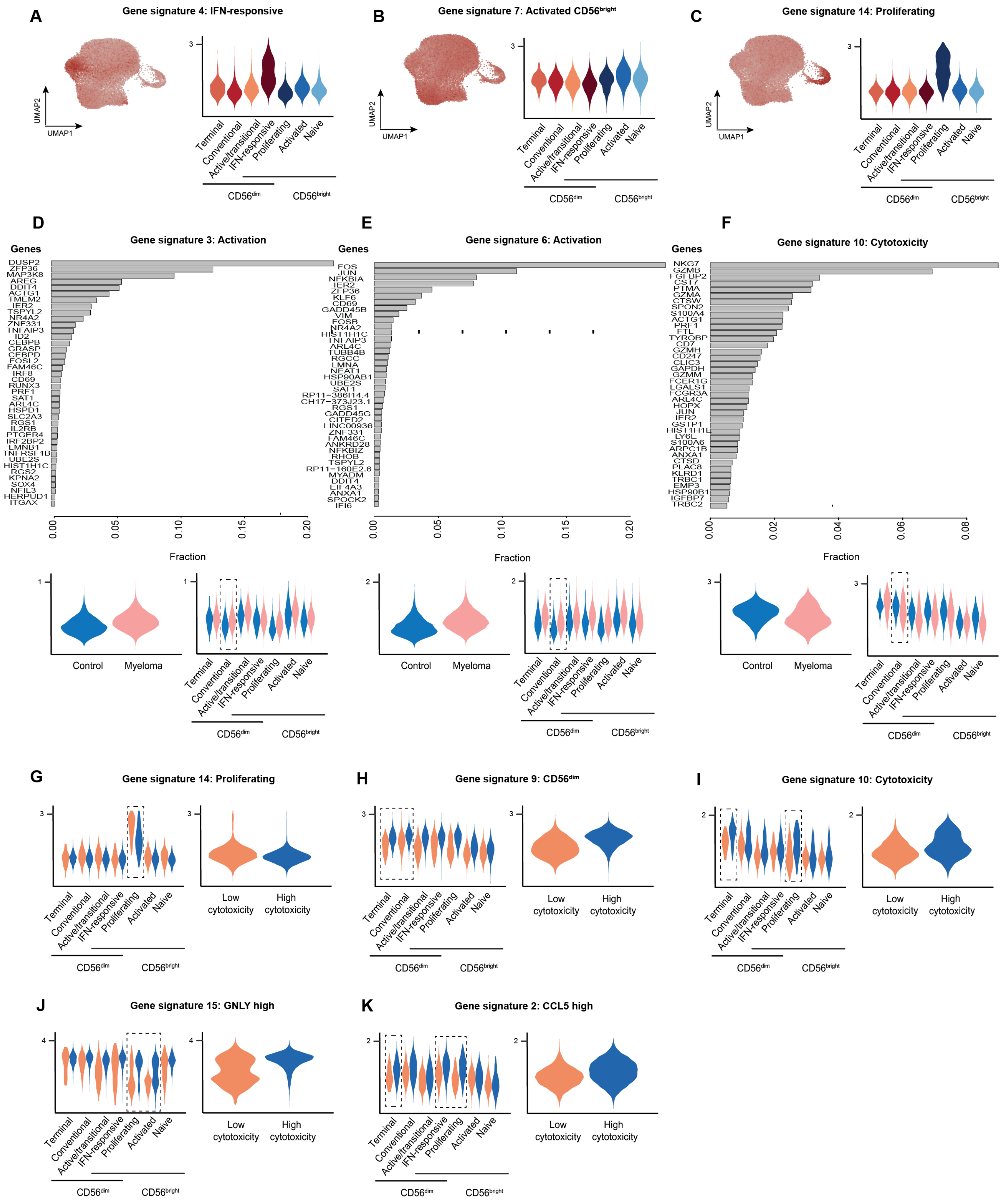
A
Gene signature 4: IFN-responsive
B
Gene signature 7: Activated CD56bright
C
Gene signature 14: Proliferating
UMAP1
UMAP2
Terminal
Conventional
Active/transitional
IFN-responsive
Proliferating
Activated
Naive
CD56dim
CD56bright
D
Gene signature 3: Activation
E
Gene signature 6: Activation
F
Gene signature 10: Cytotoxicity
Genes
DUSP2
ZFP36
MAP3K8
AREG
DDIT4
ACTG1
TMEM2
IER2
TSPYL2
NR4A2
ZNF331
TNFAIP3
ID2
CEBPB
GRASP
CEBPD
FOSL2
FAM46C
IRF8
CD69
RUNX3
PRF1
SAT1
ARL4C
HSPD1
SLC2A3
RGS1
IL2RB
PTGER4
IRF2BP2
LMNB1
TNFRSF1B
UBE2S
HIST1H1C
RGS2
KPNA2
SOX4
NFIL3
HERPUD1
ITGAX
FOS
JUN
NFKBIA
IER2
ZFP36
KLF6
CD69
GADD45B
VIM
FOSB
NR4A2
HIST1H1C
TNFAIP3
ARL4C
TUBB4B
RGCC
LMNA
NEAT1
HSP90AB1
UBE2S
SAT1
RP11−386I14.4
CH17−373J23.1
RGS1
GADD45G
CITED2
LINC00936
ZNF331
FAM46C
ANKRD28
NFKBIZ
RHOB
TSPYL2
RP11−160E2.6
MYADM
DDIT4
EIF4A3
ANXA1
SPOCK2
IFI6
NKG7
GZMB
FGFBP2
CST7
PTMA
GZMA
CTSW
SPON2
S100A4
ACTG1
PRF1
FTL
TYROBP
CD7
GZMH
CD247
CLIC3
GAPDH
GZMM
FCER1G
LGALS1
FCGR3A
ARL4C
HOPX
JUN
IER2
GSTP1
HIST1H1E
LY6E
S100A6
ARPC1B
ANXA1
CTSD
PLAC8
KLRD1
TRBC1
EMP3
HSP90B1
IGFBP7
TRBC2
Fraction
0.00
0.05
0.10
0.15
0.20
0.02
0.04
0.06
0.08
Control
Myeloma
G
Gene signature 14: Proliferating
H
Gene signature 9: CD56dim
I
Gene signature 10: Cytotoxicity
Low cytotoxicity
High cytotoxicity
J
Gene signature 15: GNLY high
K
Gene signature 2: CCL5 high

**Supplemental figure S4. Reduced frequencies of CD56$^{\text{dim}}$ NK cells in subset of NDMM patients.**

(A) Bar plot depicting scRNAseq NK cell cluster composition per individual. Relative loss of cytotoxic NK cells is labelled as 'low cytotoxicity'. (B) Representative flow cytometry gating strategy for identification of CD56$^{\text{bright}}$ and CD56$^{\text{dim}}$ NK cells in BM using the EuroFlow panel. (C) Correlation plot showing association between (B) CD56$^{\text{bright}}$ identification and (C) CD56$^{\text{dim}}$ identification using the Euroflow panel and the specifically designed panel for NK cell identification (n=7). (D) CD56$^{\text{bright}}$ and CD56$^{\text{dim}}$ levels in PB are not reflective of BM. CD56$^{\text{bright}}$ and CD56$^{\text{dim}}$ levels determined by flow cytometry using the EuroFlow panel on baseline BM aspirates of frail and unfit NDMM patients. Matched BM and PB samples were analyzed and shown in identical order of bar plots. (E) Bar plot depicting cytokine-producing CD56$^{\text{bright}}$ NK cells vs cytotoxic CD56$^{\text{dim}}$ NK cell distribution per individual in transplant-eligible treatment-naïve NDMM patients determined by flow cytometry (n=225). (F) Bar plot depicting cytokine-producing CD56$^{\text{bright}}$ NK cells vs cytotoxic CD56$^{\text{dim}}$ NK cell composition per individual in treatment-naïve frail or unfit NDMM patients determined by flow cytometry (n=44). (G) Representative flow cytometry gating strategy for CD107a degranulation in NK cells.

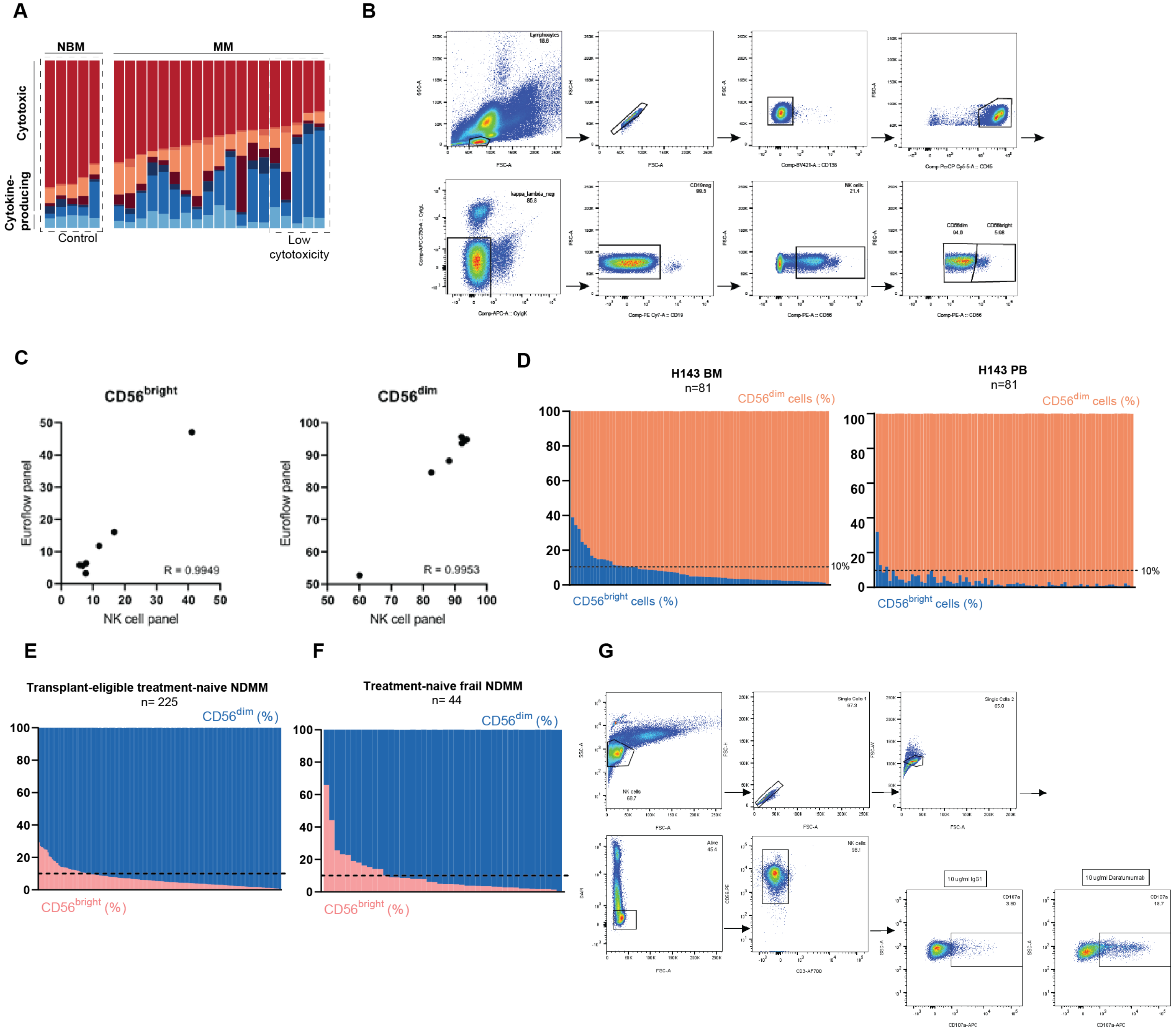

**Supplemental Figure S5. Dynamic alterations of the NK cell compartment during therapy and at relapse.** (A) Bar plot depicting BM NK cell cluster composition in MM patients post-consolidation (left) and during maintenance (right) receiving bortezomib, thalidomide and dexamethasone (VTD) with or without daratumumab (left, n=5 vs n=5; right, n=5 vs n=5). (B) Scatter plot depicting frequency of NK cell clusters per individual, split by timepoint and NK cell cluster. (C) Bar plot depicting frequency of naïve and activated CD56$^{bright}$ NK cells within the CD56$^{bright}$ compartment, split per timepoint. (D) Bar plot depicting frequency of Conventional cytotoxic CD56$^{dim}$, active/transitional CD56$^{dim}$, IFN-responsive, proliferating, and terminal NK cells within the CD56$^{dim}$ compartment, split per timepoint.

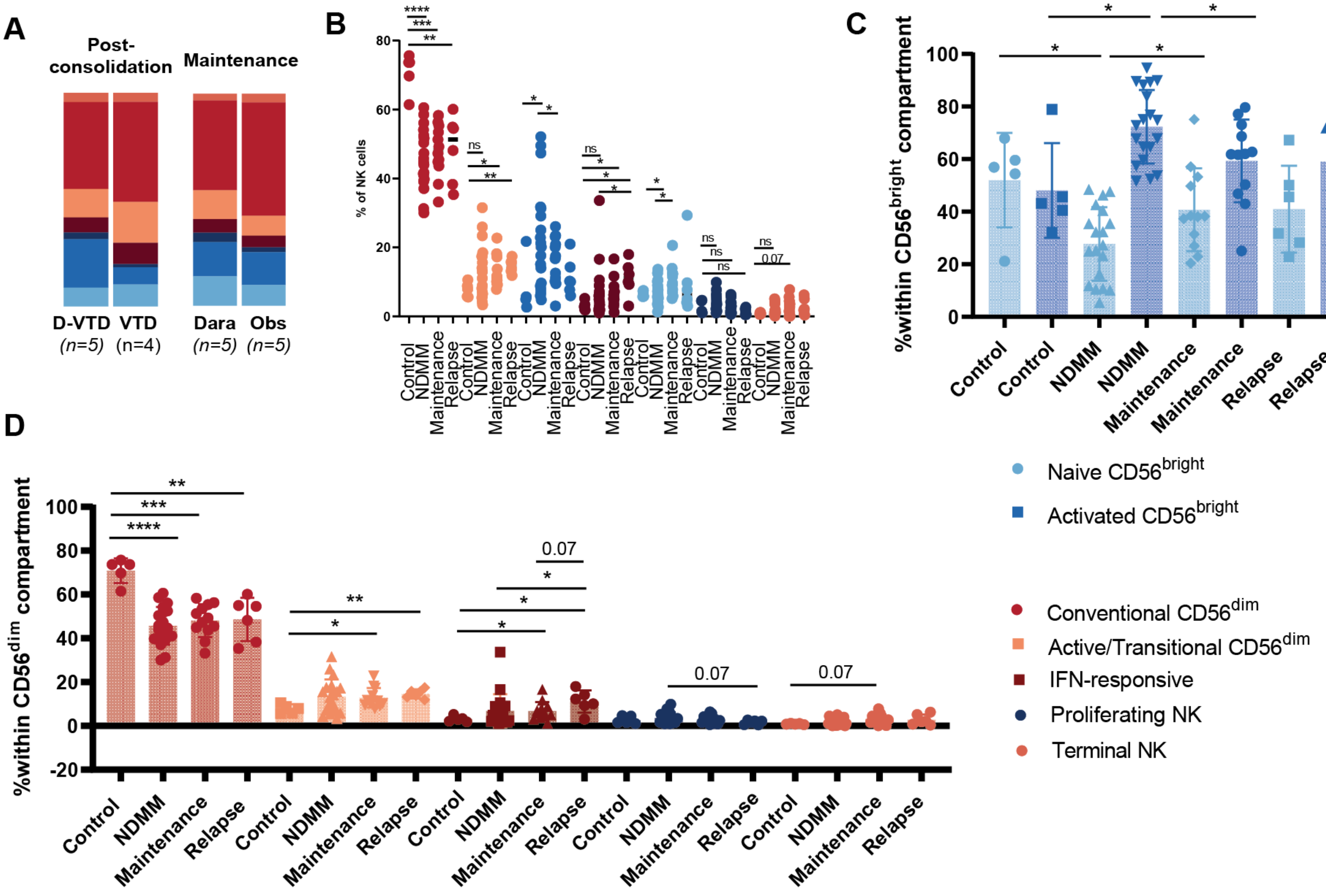

**Supplemental Figure S6. CD38⁺ expression on NK cells in the BM of NDMM patients.** A) Gating strategy to identify CD38⁺ NK cells in the BM of NDMM patients. B) The percentage of CD38 measured on CD3⁻ CD56⁺ NK cells from the BM of 7 NDMM patients. Data is presented as mean +/- standard deviation.

**A**

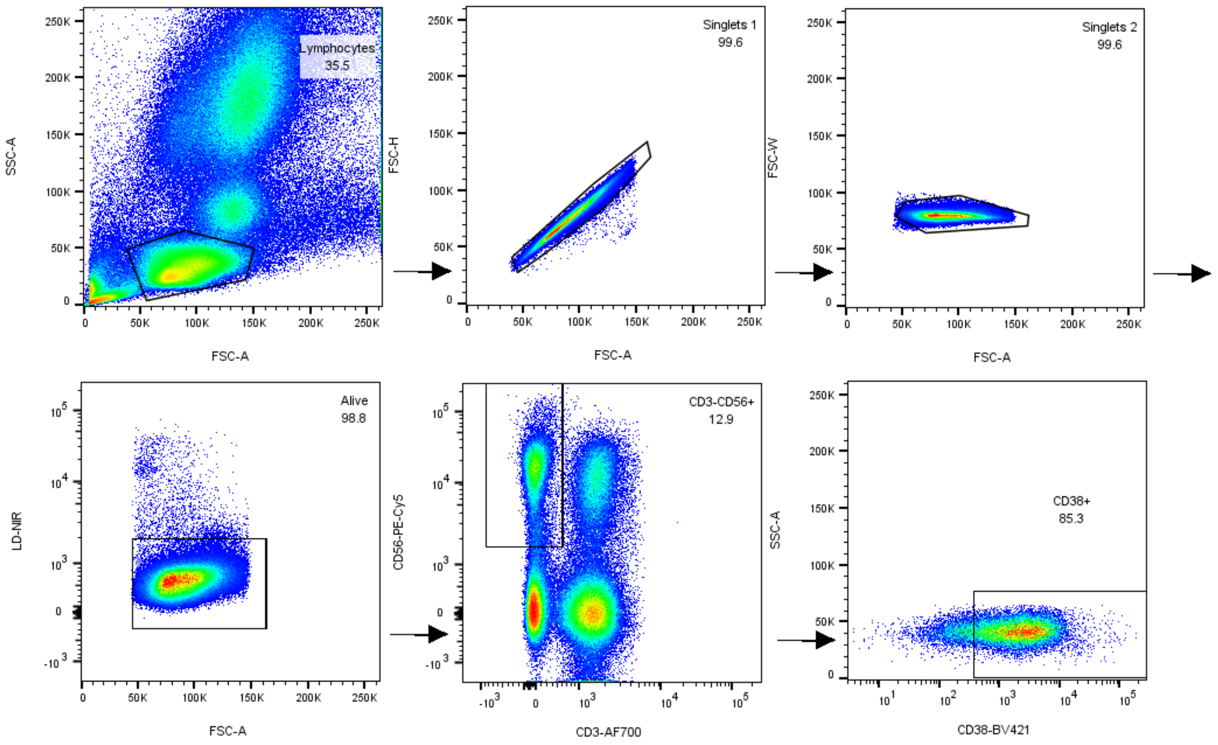


**B**

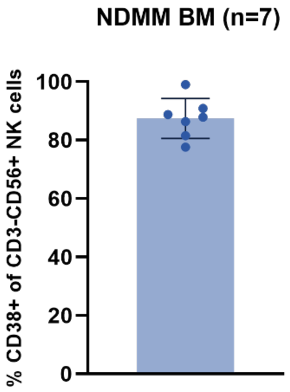

**Supplemental Figure S7. Annotation of the CD38⁻ NK cell dataset. (**A). UMAP of 9,354 human CD38⁻BM NK cells identifying three distinct clusters. Cells are color-coded according to the defined clusters. (B) Violin plots demonstrating the transcription of four NK-lineage defining markers for each cluster. (C) Feature plot depicting *NCAM1* (CD56) and *FCGR3A* (CD16) transcription among NK cell clusters.

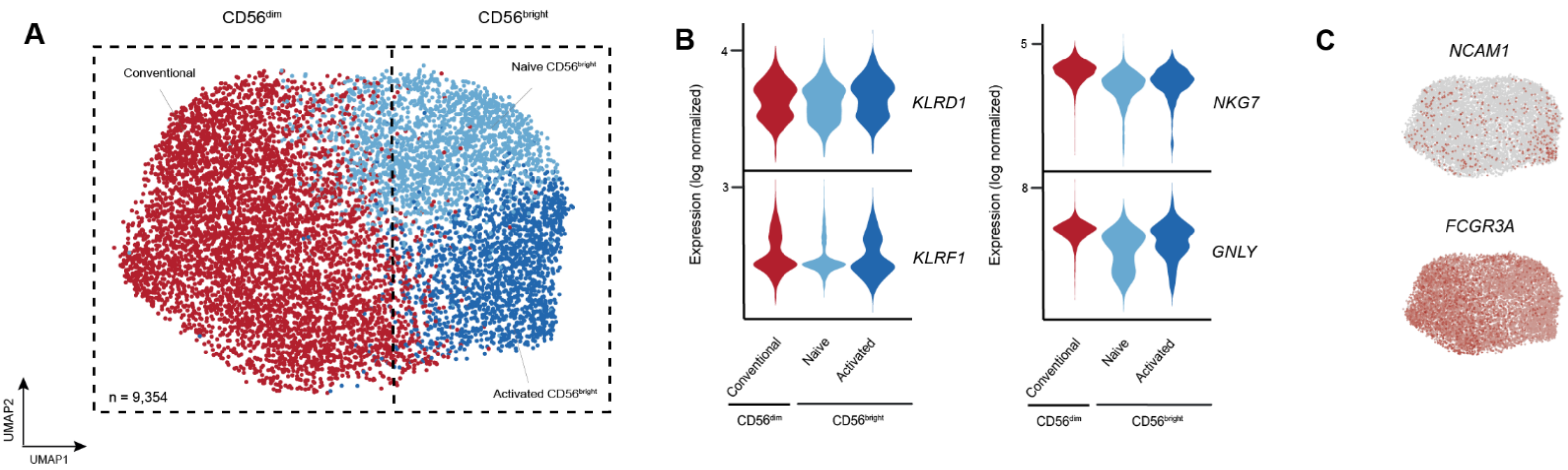

**Supplemental Figure S8. Graphical abstract.** Classically, NK cells are divided into cytotoxic CD56$^{dim}$ NK cells and cytokine-producing CD56$^{bright}$ NK cells. Normal BM consists of ~10% CD56$^{bright}$ NK cells and ~90% CD56$^{dim}$ NK cells Here, we describe the transcriptomic landscape of BM NK cells in MM. CD56$^{dim}$ NK cells are characterized by higher transcription of *GZMB*, *FCGR3A*, *KIR2DL1*, *KIR2DL3*, *PRF1, GNLY*, *LGALS1* and *B3GAT1*. CD56$^{bright}$ NK cells are characterized by higher transcription of *GZMK*, *KLRC1*, *XCL1*, *XCL2*, *CD27* and *CXCR3*. Comparison of transcriptomic profiles of BM NK cells from NDMM patients with non-cancer controls revealed reduced frequencies of cytotoxic NK cells in MM. BM NK cells in NDMM patients had reduced abundance of cytotoxicity genes (e.g. *NKG7*, *GNLY* and *PRF1*), increased abundance of inhibitory receptors *KLRB1* (CD161) and *KLRC1* (NKG2A), and decreased presence of activating receptors *NCR3* (NKp30), *CD226* (DNAM-1) and *KLRK1* (NKG2D). In approximately 20% of NDMM patients, this altered balance in cytotoxic vs cytokine-producing NK cells translated into inferior responses to therapeutic antibodies in in vitro functional assays. The reduction in cytotoxic NK cells persisted during anti-myeloma therapy and, after relapse, was accompanied by an expansion of IFN-responsive NK cells. These findings reveal substantial dysregulation of the BM NK cell compartment with potential implications for therapies dependent on ADCC. Our study offers a temporal overview of the NK cell compartment across therapy and disease progression, providing a rich resource for analyses of an important effector cell within the MM BM microenvironment.

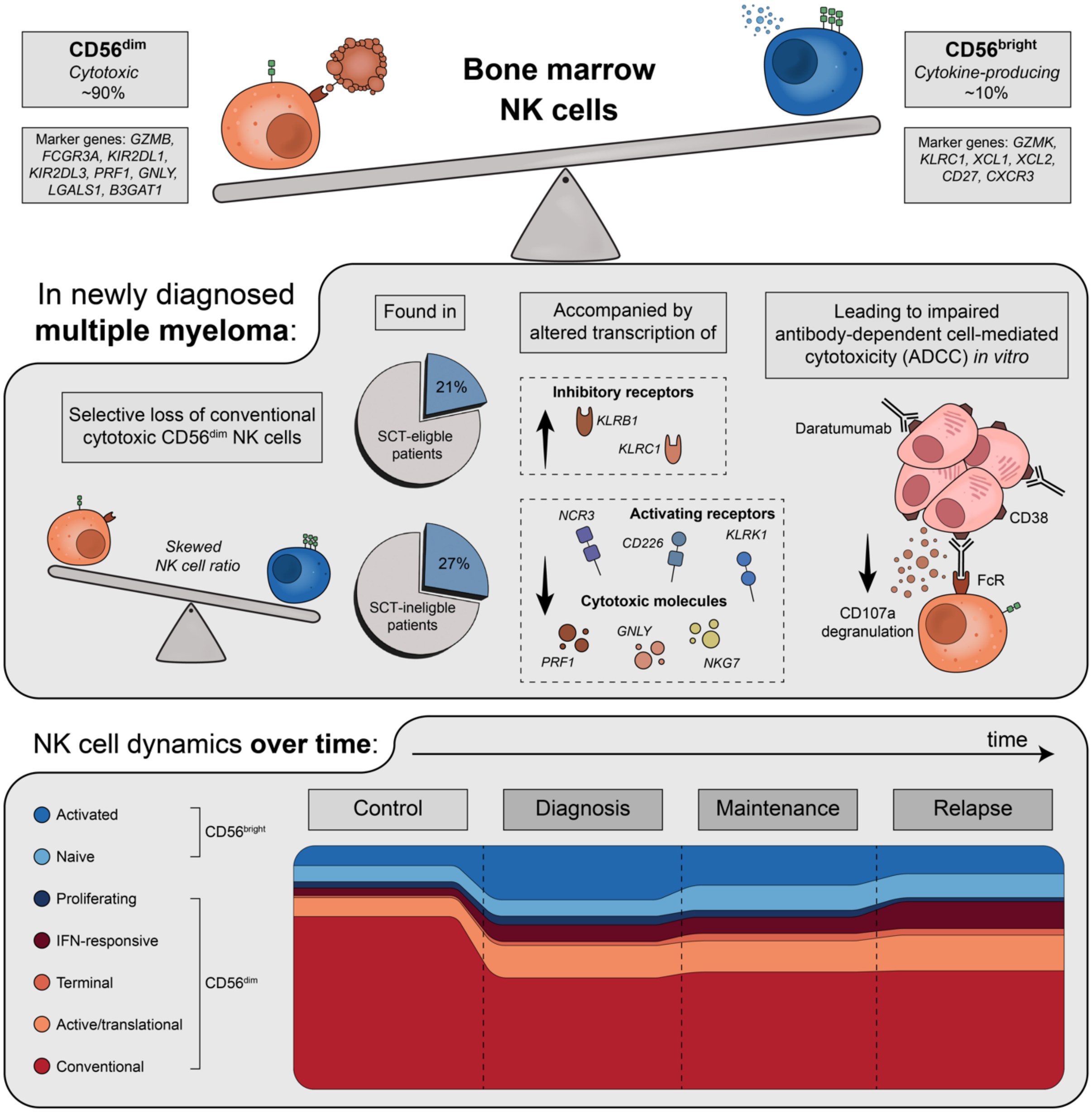
